# HDA19-mediated deacetylation of histone H3.3 lysine 27 and 36 regulates plant sensitivity to salt stress

**DOI:** 10.1101/2025.11.04.686508

**Authors:** Florian Kotnik, Minoru Ueda, Akihiro Ito, Junko Ishida, Satoshi Takahashi, Katsuyuki Sakai, Hiroshi Takagi, Julian Seidel, Takahiro Abe, Juergen Eirich, Shunji Takahashi, Dirk Schwarzer, Motoaki Seki, Iris Finkemeier

## Abstract

Plants survive extreme environments through rapid chromatin reprogramming, yet the epigenetic marks that confer stress resilience remain poorly understood. Histone deacetylase HDA19 is a key epigenetic regulator in Arabidopsis, and *hda19*-deficient mutants display tolerance to multiple abiotic stresses, including drought, heat, and salinity. Using lysine acetylome profiling, we identified a non-canonical K27/K36 di-acetylation mark on histone H3.3, among nine H3 variants, as a specific substrate of HDA19. Under salinity stress, this mark decreased in wild-type plants but increased in *hda19* mutants, while other known H3 modifications were similarly affected in both genotypes. Mimicking constitutive di-acetylation of H3.3 K27/K36 through lysine-to-glutamine substitutions promoted accumulation of stress-responsive late embryogenesis abundant (LEA) proteins and conferred salinity tolerance in seedlings, phenocopying *hda19* mutants. Furthermore, generating the *lea7-1/lea29-1/rab18-1* triple mutant abolished *hda19*-dependent salinity tolerance, confirming the LEA proteins’ role downstream of HDA19. Our findings demonstrate that H3.3 K27/K36 di-acetylation, modulated by HDA19, drives LEA protein accumulation and enables plants to withstand environmental stress, revealing a previously unknown core mechanism of plant stress resilience.

**Significance statement:** Histone acetylation is crucial for regulating chromatin states and transcription, yet the specific histone codes controlled by histone deacetylases and acetyltransferases during plant stress acclimation remain unclear. Arabidopsis mutants lacking histone deacetylase 19 show enhanced tolerance to drought, heat, and salinity. Using acetylome profiling and genetic analyses, we identify a previously unknown di-acetylation of histone H3.3 at lysines 27 and 36 as a key epigenetic switch activating late embryogenesis abundant proteins essential for salt tolerance in seedlings. This discovery reveals a novel mechanism linking histone H3.3 K27/K36 di-acetylation to stress-resilient protein accumulation, offering new insights into epigenetic control of plant stress responses and informing strategies to improve crop resilience under climate change.

## Introduction

Abiotic stressors like salt stress pose significant threats to crop yields. Hence, the elucidation of the molecular mechanisms underlying plant stress responses is crucial for developing resilient crops. Histone deacetylase 19 (HDA19), also known as HDA1 or HD1, plays a key role in regulating gene expression during plant development and in response to stress (1-4). However, the mode of action of HDA19 in driving epigenetic regulatory machinery in response to stress remains unclear.

Within the nucleus of eukaryotic cells, core histone proteins (H2A, H2B, H3 and H4) wrap DNA into higher order structures called nucleosomes, enabling sophisticated regulation of nuclear gene expression and protecting DNA from physical damage. Multiple post-translational modifications (PTMs) can occur on the N-terminal tails of histones and play a crucial role in regulating chromatin structure and, consequently, gene expression. The most important histone PTMs include lysine acetylation, various other acylations, lysine and arginine methylation, lysine ubiquitination as well as serine and threonine phosphorylation. Based on the interplay of these modifications, which are essential for chromatin structure and gene activity regulation, a histone code theory was proposed (5-8). Previous studies have established a positive correlation between levels of lysine acetylation at N-terminal histone H3 sites and active gene expression (9, 10). The interplay between histone acetyltransferases (HATs) and HDACs acts as a dynamic regulatory mechanism, with these enzymes functioning as writer and eraser proteins that modulate the acetylation status at distinct histone sites (11). Reader proteins, including bromodomain-containing proteins, specifically recognize acetylated sites and recruit transcriptional complexes to target genes, thereby determining their activation state. Notably, various histone H3 variants, characterized by subtle alterations in their tail domains, have been shown to play unique roles in regulating gene expression and plant development. For instance, histone H3.3 is predominantly associated with transcriptionally active regions during Arabidopsis development, whereas the H3.1 variant is more commonly found in inactive heterochromatic regions (12).

In Arabidopsis, the HDAC family comprises three classes: HDA1/RPD3-type, sirtuin, and HD-tuin subfamilies, consisting of 18 genes. Each HDAC has diversified its function and plays pivotal roles in various biological processes (13). As a member of the RPD3-type subfamily of class I histone deacetylases, HDA19, together with HDA6 and HDA9, is involved in the epigenetic regulation of gene expression. HDA19 deficiency leads to a high tolerance towards salt, dehydration, and heat stresses with slower development and reduced fertility in Arabidopsis Col-0 and rice (13-16). Despite growing recognition of the importance of HDA19 and its homologue in rice in stress responses (17), its substrates remain unclear. Most studies have focused on its role in regulating specific histone marks, such as H3K9ac, H3K9K14ac, and H3K9K18ac, which are commonly conserved among eukaryotes including yeast, plants, and mammals (18).

Recent mass spectrometry-based (MS) lysine acetylome studies in plants and mammalian cell culture using the RPD3-class inhibitor apicidin have comprehensively revealed significant changes in the nuclear acetylome when the activity of this class of deacetylases is modulated, affecting both histone and non-histone proteins (19, 20). The diversification of primary structures of plant histones (H2A; 6 variants and 13 genes, H2B; 11 variants and 11 genes, H3; 9 variants and 15 genes in Arabidopsis), except for H4 (21), and the emergence of plant specific accessory proteins like angiosperm-specific HDA19-interacting homologous proteins (HDPIs) (22), highlight the need for a comprehensive dissection of *hda19* mutants to uncover acetylation marks through which HDA19 dominantly regulates stress responses. In this work, we found that di-acetylation of H3.3 at K27/K36 mediates salt tolerance, whereas the absence of HDA19 has only a minor effect on canonical acetylation marks, including H3K9ac, H3K9K14ac, and H3K9K18ac. Furthermore, lysine-to-glutamine (K/Q) substitutions, which mimic the acetylated states at this novel di-acetylation site, induce the expression of late embryogenesis abundant (LEA) proteins, thereby conferring salt stress tolerance comparable to that observed in *hda19* mutants.

## Results

### *Rfhda19*, a novel *hda19* allele with reverted fertility and maintained stress tolerance

Our previous work showed that the *hda19-3* mutant, which is deficient in HDA19, exhibits a severe defective phenotype in reproductive organ-formation and fertility, as well as a salt and drought stress tolerance phenotype. In this study, we aim to identify the *in vivo* HDA19 substrates by mass spectrometry-based acetylome profiling to elucidate the mechanism by which *hda19* mutants acquire their tolerance. The slow growth of the *hda19-3* mutant made it challenging to directly compare them with Col-0 wild-type (WT) plants. To address this issue, we screened for additional *hda19* mutant alleles that retained their stress tolerance but had a less severe growth defect. For this screen we used the *hda19-7* line, which was selected from a CRISPR-Cas mediated genome-editing experiment as described previously (16). *hda19-7* contains an additional G insertion 79 nucleotides downstream of the start codon within *HDA19* coding sequence. Among the progeny of this line, we identified a revertant in fertility, designated *Rfhda19* for this reason. This *Rfhda19* mutant contains an additional insertion of plasmid-like foreign DNA into the genomic region around the translational initiation site of HDA19, resulting in the deletion of 57 nucleotides from the HDA19 coding sequence, which leads to a modified N-terminus of the protein (*SI Appendix*, Fig. S1*A*).

*Rfhda19* retains similar stress resistance to *hda19-3* and exhibits partial recovery to semi-fertility from almost sterile phenotypes, enabling better growth and allowing for a more suitable comparison to WT plants (Fig. 1*A*, *SI Appendix*, Fig. S1, *B* to *D*, Fig. S2*A*). To confirm that the phenotype is caused by the lack of active HDA19, the genomic DNA encompassing the coding sequence and proximal regulatory elements of *HDA19* was introduced into *Rfhda19*, and the resultant T_3_ plants displayed decreased tolerance to salinity stress as WT plants (*SI Appendix*, Fig. S1 *E* and *F*). To determine if the *Rfhda19* line also lacks an active HDA19 enzyme, we performed a pull-down analysis using the HDAC-trap mini-AsuHd, which enriches catalytically active HDACs by hydroxamic acid grafted onto a lysine-like scaffold (23). While HDA19 was highly enriched on mini-AsuHd from WT extracts, it was absent in *Rfhda19* mutants (*SI Appendix*, Fig. S3 and Dataset S1) due to its impaired activity.

**Figure 1.**
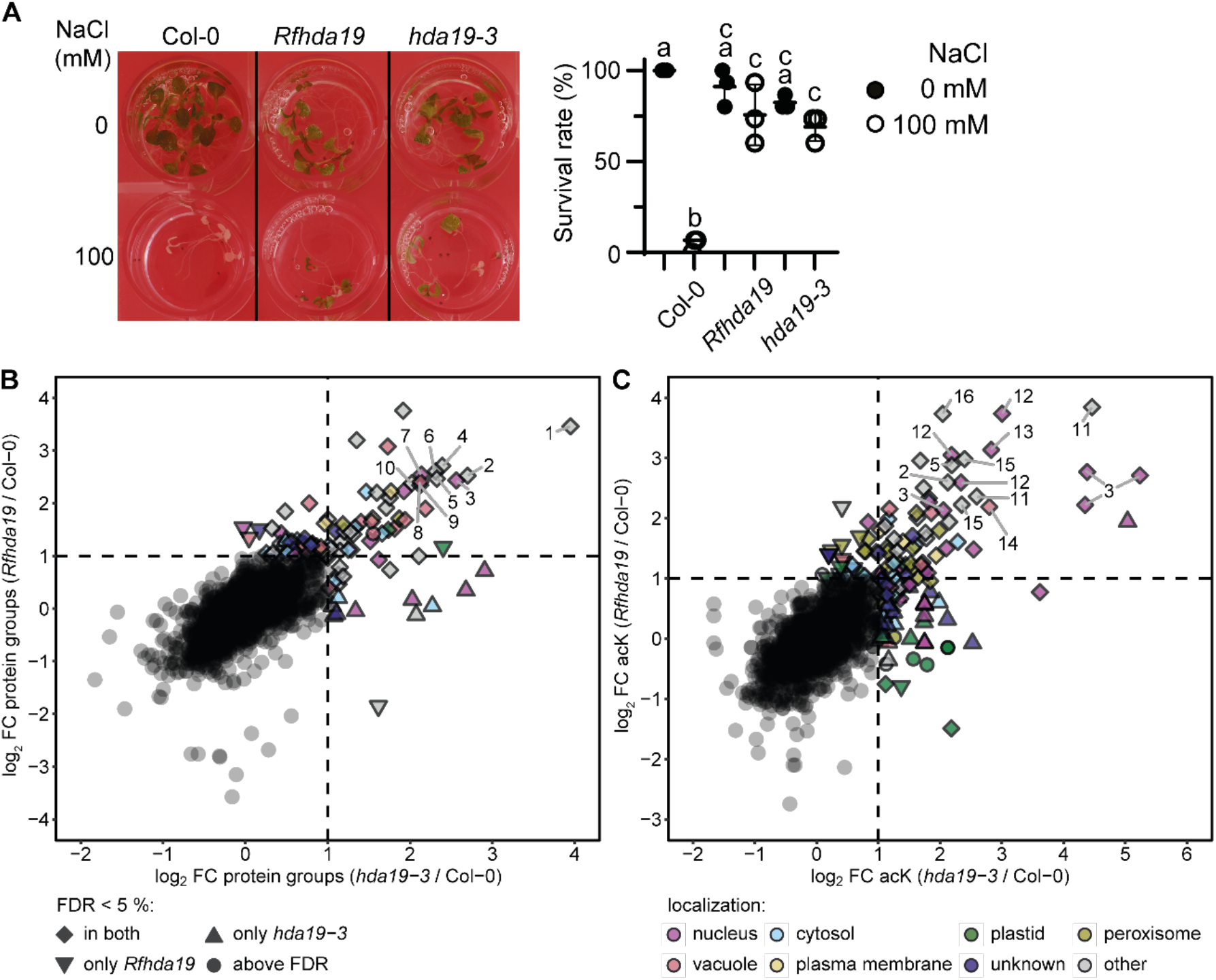
Identification of lysine-acetylated HDA19 substrates involved in plant stress tolerance.(*A*) *Rfhda19*, a revertant in fertility of the *hda19-7* mutant, exhibited similar tolerance to salinity stress (100 mM NaCl) as *hda19-3*. Multiple comparisons of survival rate were performed using one-way ANOVA, with *P* < 0.05 considered significant (n=3). Inside diameter of the well is 15.4 mm. (*B* and *C*) Scatter plots comparing (*B*) protein groups and (*C*) lysine acetylation (acK) site abundance changes in *hda19-3* vs WT and *Rfhda19* vs WT. The original data can be found in Dataset S2 and S3. Protein names or symbols of numbered spots are displayed in *SI Appendix*, Table S1.

### Histone H3.3 and seed-specific proteins are hyperacetylated *in hda19*

To identify the *in vivo* protein substrates of HDA19, we analyzed the proteomes and acetylomes of five-day-old Arabidopsis seedlings from the two different lines lacking HDA19 (*hda19-3* and *Rfhda19)* compared to WT plants using a quantitative liquid chromatography mass spectrometry-based (LC-MS) proteomic approach. In total, 4989 and 4428 protein groups were quantified in at least two biological replicates in the *Rfhda19* and *hda19-3* versus WT comparisons, respectively. Moreover, 3726 and 3318 lysine acetylation (acK) sites were quantified on 1669 and 1465 unique protein groups, respectively (*SI Appendix*, Fig. S4 and S5, Dataset S2 and S3). The majority of protein groups and 1912 lysine acetylation (acK) sites were common to both *hda19* deficient lines in these datasets (*SI Appendix*, Fig. S6). To reveal whether similar changes in protein and acK site abundance changes occur in both *hda19* mutant lines, we directly compared the regulation patterns between the two lines. Scatter plots illustrate a high degree of similarity between the two *hda19* mutants relative to WT, with 44 protein groups and 63 acK sites on 36 protein groups exhibiting strong increases in abundance in both lines (log_2_ FC > 1) (Fig. 1,*B* and *C*). Given this high degree of correlation of the proteome response in both lines, we combined them into a single group (*hda19*) and considered protein groups and acK sites quantified in six out of eight replicates for further statistical analysis. This combined analysis revealed 94 acK sites on 49 protein groups with significantly and more than two-fold increased hyperacetylation (log_2_ FC > 1, FDR < 5%) (Fig. 2,*A* and *B*, Dataset S4). The strongest increase in lysine acetylation (>16-fold) was observed for the seed storage protein CRUCIFERIN 4 (CRU4) and LEA29 (*SI Appendix*, Fig. S7). Only 11 of the 49 hyperacetylated proteins (AHL14, Ankyrin repeat family protein (AT2G26210), DNA-binding storekeeper protein-related transcriptional regulator (AT4G00390), H3.3, LEA7, LEA25, LEA29, MAC7, PARP3, PWO1, SCS2B) are known to localize to the nucleus, and one (RAB18) to the nucleus and plasma membrane, according to SUBA5 (Fig. 2*C*). A sequence logo generated from the 94 hyperacetylated sites revealed a putative substrate motif for HDA19, characterized by lysine residues at positions −13, −2, and +11, and alanine residues at positions −3 and +1, and aspartic acid at position −1 surrounding the actual acetylation site (Fig. 2*D*). Functional enrichment analysis of the hyperacetylated proteins showed that the GO Component term “protein storage vacuole”, the GO Function term “nutrient reservoir activity”, and the UniProt Keyword “seed storage protein” were the three terms most enriched terms (Fig. 2*E*). Comparison of proteome and acK site abundance changes revealed a high correlation, with 24 protein groups exhibiting similar increases in acetylation and protein abundance in *hda19* compared to WT (Fig. 2*F*). However, 24 acK sites on 19 proteins showed an at least 2-fold stronger increase in lysine acetylation than in protein abundance. Among these 19 proteins, the histone protein H3.3 with two acetylation sites (K27 and K36) and multiple LEA proteins with various acK sites (LEA25 K300; LEA29 K38, K82, K111; LEA48 K153; RAB18/LEA51 K180) were identified.

**Figure 2.**
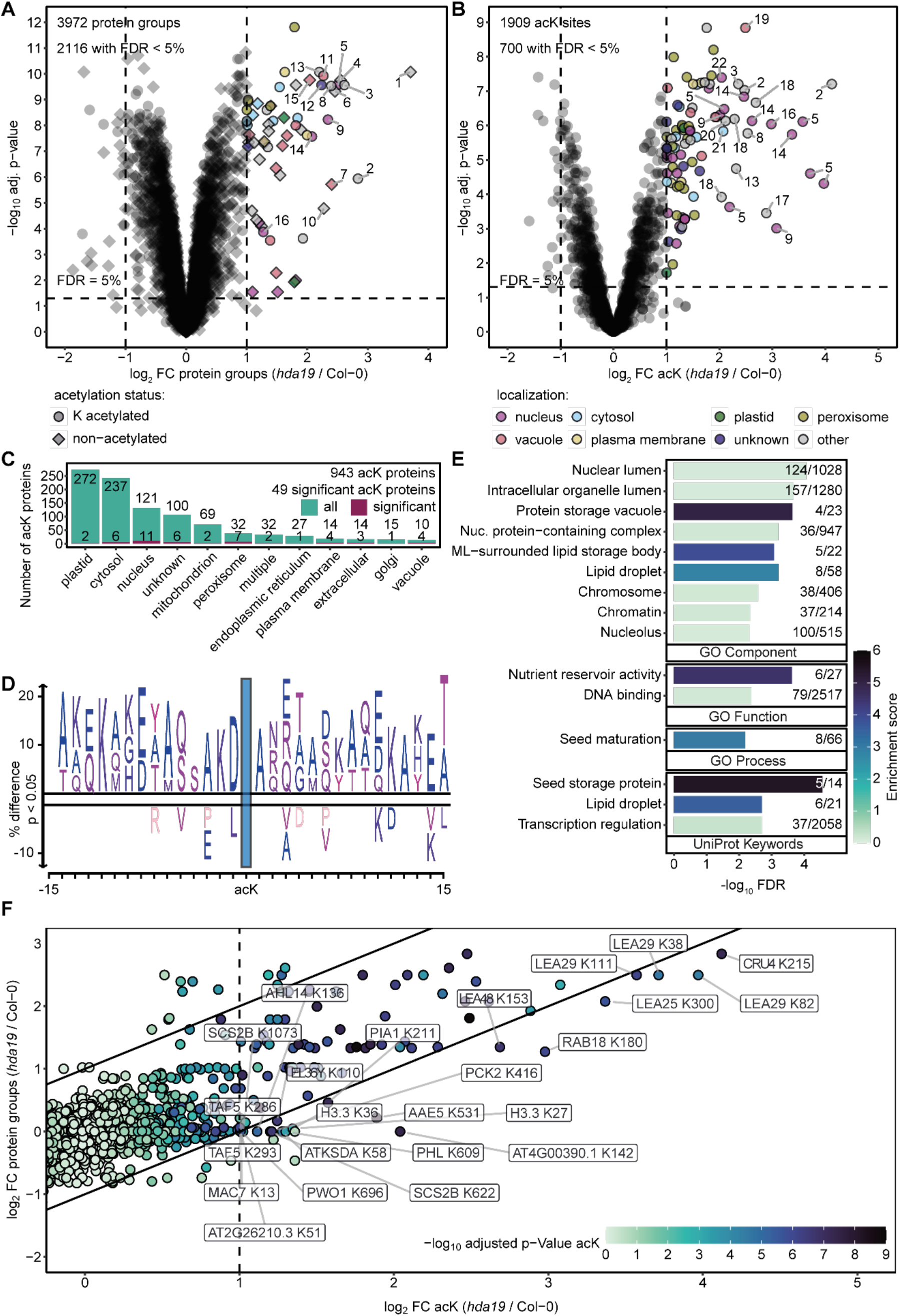
Identification of lysine-acetylated HDA19 substrates involved in plant stress tolerance. (*A* and *B*) Volcano plots showing (*A*) protein groups and (*B*) lysine acetylation (acK) site log_2_ fold abundance changes in *hda19-3* compared to Col-0 WT, along with −log_10_(adjusted p-values) from *limma* analysis (74). The p-values were adjusted using a Benjamini-Hochberg correction. Data from *hda19-3, Rfhda19* and WT were processed and analysed as one *hda19* group, and only acK sites and protein groups quantified in at least 6 of 8 biological replicates were included in the analysis. Log_2_ fold changes (FC) of +/-1 and false discovery rates (FDR) of < 5 % are indicated by dashed lines. Significantly altered proteins and acK sites (log_2_ FC > 1, FDR < 5 %) are colored according to their SUBA5 localization (75). AcK sites and proteins with a log_2_ FC > 2 are numbered. Protein numbers with symbols or identifiers from 1 to 22: 1. AT2G05580, 2. CRU4, 3. CRU3, 4. PAP85, 5. LEA29, 6. AT4G36700, 7. LEA18, 8. CRU1, 9. LEA7, 10. CRU2, 11. OLEO2, 12. OLEO4, 13. LEA30, 14. LEA25, 15. CLO1, 16. RAB18, 17. SESA1, 18. LEA48, 19. OLEO1, 20. LEA42, 21. LEA36, 22. AT4G00390. The original data is available in DataS4. (*C*) Subcellular localization of all quantified acetylated proteins (green), and those which are significantly regulated in *hda19* vs WT (purple) with a log_2_ fold-change (FC) acK > 1, FDR < 0.05. (*D*) Sequence logo of significantly upregulated acK sites in *hda19* (log_2_ FC > 1, FDR < 0.05). The blue box indicates the position of the acetylated lysine residue. (*E*) Functional enrichment analysis of all quantified acetylated proteins in *hda19* vs WT. The bars represent the −log_10_(FDR) for each term. The color of each bar corresponds to the enrichment score for each term, calculated by STRING. The numbers adjacent to the bars indicate the number of proteins found and the total number for each term. The analysis included GO component, GO function, GO process, and UniProt keywords. Only terms showing an increase in abundance are displayed. (*F*) Comparison of protein and acK log_2_-fold changes in *hda19* (*hda19-3* + *Rfhda19*). Nineteen proteins exhibited a significant increase in acetylation at 24 acK sites, exceeding their respective protein abundance changes. The lysine (K) positions of the histone sites were adjusted to account for the removal of the initiator methionine. The original data can be found in Dataset S4.

### Histone H3K27/K36 di-acetylation is indispensable for salinity tolerance

Gene expression is heavily influenced by histone modifications, with even minor variations in histone sequences giving rise to distinct regulatory functions. In Arabidopsis, 50 genes encode 37 different histone proteins (24). We quantified 32 of those histone proteins at either the protein or acetylation level (Dataset S4). Due to the high lysine content of histone tails, tryptic peptides often carry more than one acK site. This is particularly important, as H3.3 hyperacetylation was only observed on peptides carrying di-acetylation at both K27 and K36, while peptides showing mono-acetylation only at K36 were slightly downregulated (Fig. 3*A*). Since the differences in amino acid sequences between the histone variants are relatively small, distinguishing between them is not always possible, especially for acK sites where the corresponding amino acid changes may not reside on the same tryptic peptide. In the case of H3.3, a single amino acid exchange from alanine to threonine between H3.1 and H3.3 can be found at position 31 (24), which is distinguishable in the MS/MS spectra of the di-acetylated peptide (Fig. 3*B*). H3.3 is often enriched at euchromatin regions and associated with active gene expression, whereas H3.1 is enriched in heterochromatin and associated with gene silencing (25). Both acetylome and immunoblotting analyses detected a significant increase in di-acetylation levels of lysines (K) 27 and 36 of histone H3.3 (H3.3K27/K36ac) in *hda19* compared to WT plants, although this increase was less pronounced in *Rfhda19* (Fig. 2*F*, Fig. 3,*A* and *C*). The difference between *hda19-3* and *Rfhda19* may be attributed to the fact that a truncated, non-active HDA19 protein might be generated in *Rfhda19*, although this was not detectable by LC-MS/MS analysis. Additionally, the difference could be caused by the slightly enhanced developmental speed of *Rfhda19* compared to *hda19-*3, particularly given that H3.3 enrichment patterns change during early development (26). Plants treated with trichostatin A (TSA), a pan-HDAC inhibitor, showed a 2-fold increase of acetylation at H3.3K27/K36ac compared to control conditions in the immunoblot analysis (Fig. 3*C*). Similarly, a previous MS-based analysis of apicidin treated plants showed a nearly 6-fold increase in di-acetylation of H3.3K27/K36ac (19). Higher acetylation levels of K27/K36 after inhibitor treatments may be explained by additional inhibition of HDA6, which is thought to work redundantly with HDA19 (27). In contrast to *Rfhda19* treated with salt, WT plants showed slightly decreased di-acetylation of K27 and K26 under the same conditions (Fig. 3*A*). This is consistent with a previously observed up-regulation of HDA19 activity upon salt stress in WT plants (28). Here we detected increased lysine acetylation on H3.3 only in the absence of HDA19 (Fig. 3*A*), indicating that HDA19 can distinguish between histone variants to regulate gene expression.

**Figure 3.**
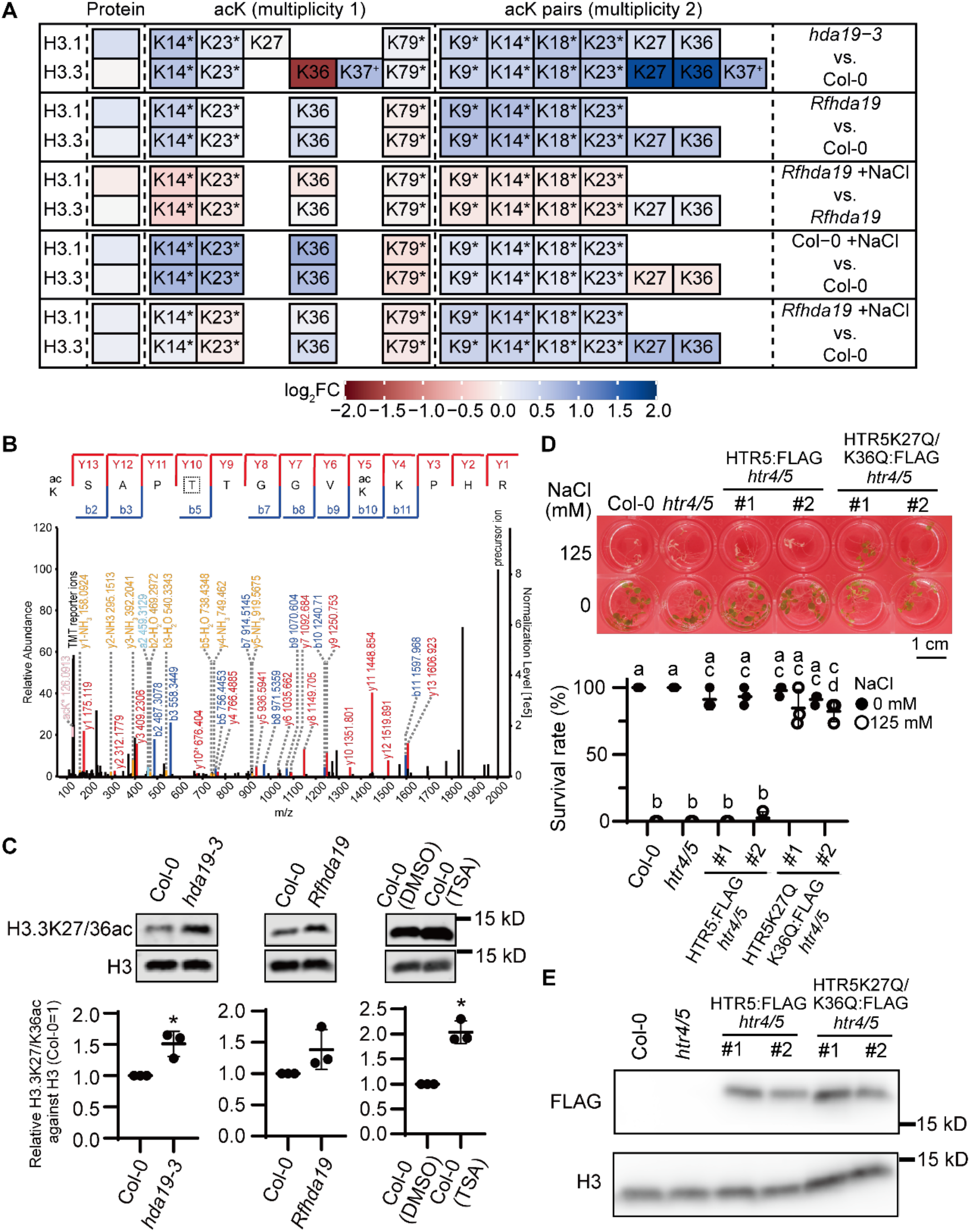
Requirement of H3.3K27/K36 di-acetylation for enhanced tolerance to salinity stress. (*A*) Protein and acK abundance of the two main histone 3 variants, H3.1 and H3.3, in *hda19-3, Rfhda19, Rfhda19* + NaCl, and WT + NaCl, were quantified in at least 2 out of 4 biological replicates. Colors indicate the respective log_2_ fold change (FC) values. Lysine positions have been adjusted to reflect the cleavage of the initiator methionine. The histone proteins can carry multiple acetylations and the number of distinct modified versions of the same peptide sequence that exist or are detected is therefore called multiplicity. Hence, lysine acetylation (acK) sites are grouped by their respective multiplicity, as determined by MaxQuant. Stars next to the lysine (K) position indicate sites that could belong to both histone variants. Di-acetylated (multiplicity 2) peptides of H3.1 and H3.3 can be found in *SI Appendix*, Table S2. For acK sites marked with a +, we were not able to clearly distinguish the modification sites K36 and K37 for all peptides, since these residues are positioned next to each other in the peptide sequence. Distinctive fragment ions could be identified to unambiguously localize K36ac, but not K37ac. The localization probability of K37ac is around 0.5. (*B*) MS/MS spectra, as processed by MaxQuant, show an H3.3 peptide carrying K27 and K36 di-acetylation. The amino acid at position 31, which differentiates H3.3 from H3.1, is marked by a dashed box. (*C*) Differences in histone H3 acetylation levels in the *hda19-3, Rfhda19*, and plants treated with trichostatin A (TSA) were evaluated. Quantification of relative acetylation levels was performed using Student’s *t*-test (n=3, *P* < 0.05). (*D*) Expression of FLAG-tagged HTR5K27Q/K36Q acetylation mimicking variant in *htr4/5* enhanced tolerance to salinity stress (125 mM NaCl). Multiple comparisons of survival rate were performed using one-way ANOVA, with *P* < 0.05 considered significant (n=3). (*E*) Immunoblotting of the FLAG-tagged HTR5 and FLAG-tagged HTR5K27Q/K36Q is shown. The full blot is presented in *SI Appendix*, Fig. S1.

To confirm the role of H3.3K27/K36ac in response to salinity stress, we introduced two K to Q mutations at the 27 and 36 sites of histone HTR5, a variant of H3.3 in Arabidopsis. The replacement of lysine residues with glutamine (K→Q) is considered to mimic the acetylation mark due to the permanent abolishment of the positive charge (29). Three different genes, HTR4/5/8, encode H3.3 in Arabidopsis (24). Expression of HTR5:FLAG complemented the *htr4/5/8* mutant derived from *htr4/5*(-/-) *htr8*(+/-) progeny (*SI Appendix*, Fig. S8), indicating that FLAG-tagged HTR5 is functional, whereas constitutive expression of K27Q/K36Q in all three H3.3 variants appears lethal in Arabidopsis, as HTR5K27Q/K36Q:FLAG was detected only in an *htr4/5* background but not in *htr4/5/8* progeny. Plants expressing HTR5K27Q/K36Q:FLAG under the control of the gene-specific promoter and terminator showed salinity stress tolerance (Fig. 3,*D* and *E, SI Appendix*, Fig. S2*B*). These data indicate that H3.3K27/K36ac regulates the response to salinity stress in Arabidopsis.

### Identification of HDA19 interaction partners in Arabidopsis seedlings

Previous studies have identified several proteins that interact with HDA19 in chromatin-modifying complexes. To further investigate the interactions of HDA19, we performed a quantitative pull-down analysis of MycHDA19 expressed in *hda19-6* seedlings compared to WT plants via LC-MS. In total, 203 protein groups were quantified in at least two out of four replicates, and 22 of those protein groups were significantly enriched (log_2_ fold-change > 1 and adjusted p-values (FDR) < 0.05) in MycHDA19 compared to WT, including HDA19 itself (Fig. 4, Dataset S5). Among these 22 proteins, nine localize to the nucleus and five to the cytosol, suggesting that HDA19 might also function in more than one compartment. We detected multiple members of the known HDA19-SIN3 complex, including HISTONE DEACETYLASE COMPLEX 1 (HDC1), MSI1, SIN3-like (SNL) proteins 1-6 (14, 30-32). Recently, three new angiosperm specific members of the HDA19-SIN3 complex called HDIPs were identified (22). HDIP3 co-purified with MycHDA19 in this study, along with several previously unreported proteins (Fig. 4), including the small GTP-binding protein RAN1, which regulates nuclear-cytoplasmic transport (33), four ribosomal proteins, and the dynamin-related protein 2 family protein important for gametophyte development and biotic stress resistance (34, 35). These HDA19 interacting proteins are likely to play a crucial role in regulating stress responses through the modification of H3.3K27/K36ac.

**Figure 4.**
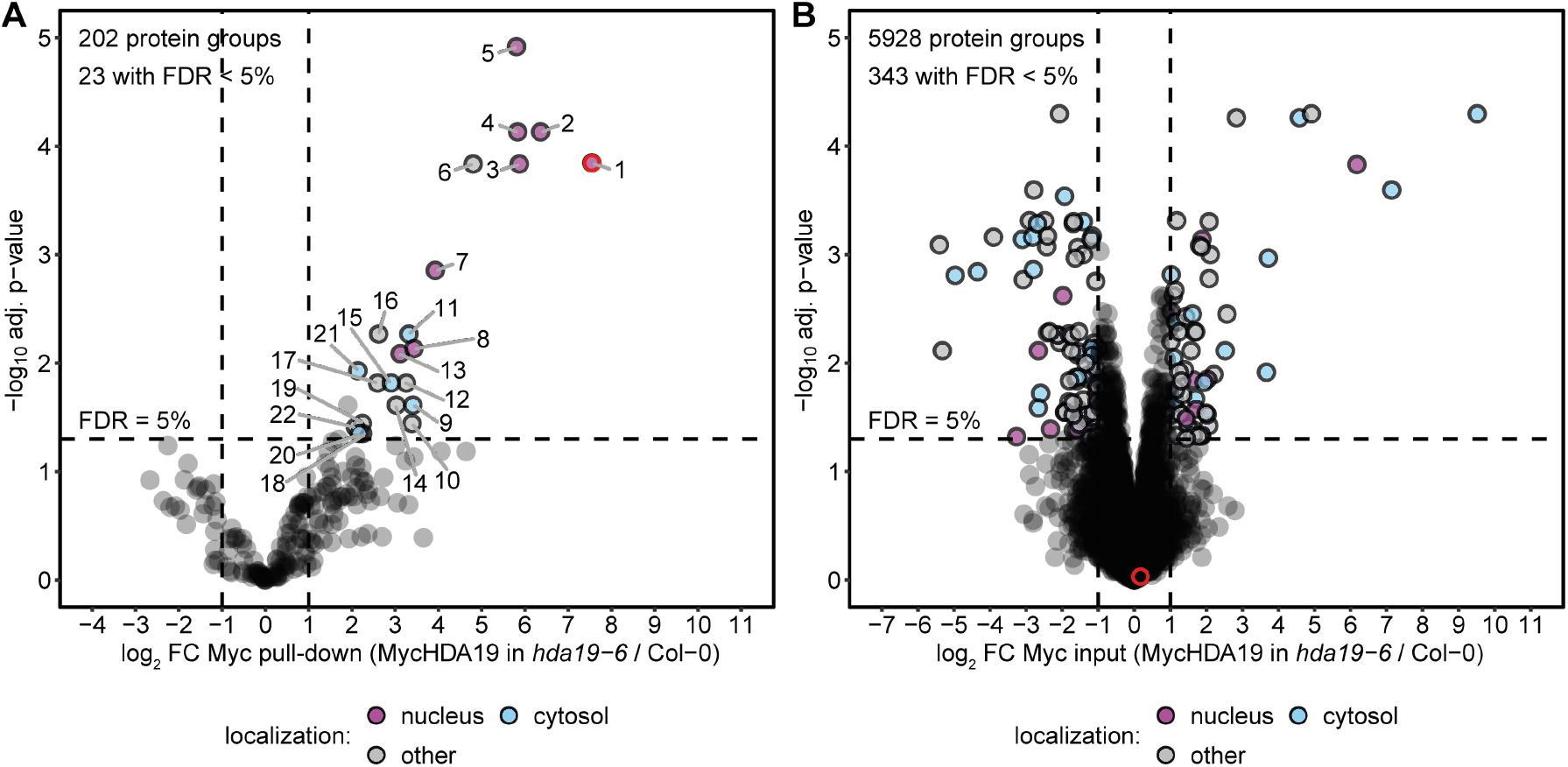
Identification of HDA19 interacting proteins. Volcano plots showing (*A*) enriched proteins in Myc pulldown and (*B*) overall protein abundance changes (input) in MycHDA19 (in *hda19-6*) compared to Col-0 WT, along with −log_10_(adjusted p-values) from *limma* analysis (74). The p-values were adjusted with Benjamini-Hochberg correction. Protein groups only quantified in at least 2 of 4 biological replicates were included in the analysis. Log_2_ fold changes of +/-1 and FDR of 5 % are indicated by dashed lines. Significantly enriched proteins (log_2_ FC > 1, FDR < 0.05) are colored according to their SUBA5 localization (75). Significantly enriched proteins with a log_2_ FC > 2 are numbered. Protein numbers with symbols or identifiers from 1 to 22: 1. HDA19, 2. HDC1, 3. ES25Y, 4. MSI1, 5. SNL5, 6. SNL4, 7. SNL6, 8. SNL2, 9. RPL5A, 10. AT5G36790, 11. CCT7, 12. HDIP3, 13. HB1, 14. FdC1, 15. RAN-1, 16. APOK3, 17. ICL, 18. SNL1, 19. CRU2, 20. RPS18C, 21. RPS26e, 22. LACS6. HDA19 is highlighted by a red circle in A and B. The original data can be found in Data S5.

### Requirement of LEA proteins in enhancing salinity stress tolerance in *hda19* mutants

LEA proteins play crucial roles in protecting plants against environmental stresses, such as desiccation, drought and salinity. Despite their importance, the regulation of LEA proteins is not yet fully understood. In total, 22 out of 51 LEA proteins were quantified at either the protein or acK level (Fig. 5*A*). These proteins belong to different subfamilies, including LEA_1, LEA_2, LEA_3, LEA_4, dehydrins, and PvLEA18. LEA_4 family is the largest of these, comprising 19 LEA proteins, of which nine were found in this dataset (36). Here we report that the protein abundance of 11 LEA proteins and 33 acK sites of seven of these proteins were strongly upregulated in *hda19* mutants (Dataset S4). The strongest induction was primarily observed for the LEA_4 family members, with many proteins showing significant increases in both protein and acK site abundance. The highest increase was observed for LEA29 acK sites K38, K82, and K111, which showed an over 11-fold increase in the *hda19* mutants compared to WT plants (Dataset S4). LEA7, a paralog of LEA29 with preserved nucleotide similarity, was duplicated as a result of a whole-genome duplication event (36) and also exhibited increased protein abundance and acK levels in *hda19* mutants. In addition to the LEA_4 family proteins, the acetylation of the dehydrin RAB18/LEA51 was also strongly increased at K180 (Fig. 5*A*). Both LEA7 and LEA29 have been reported to be induced under salt and osmotic stress, and during treatment with the plant hormone abscisic acid (ABA) (37). RAB18 is also induced in response to salinity, ABA, and other environmental stresses (36, 38). To test whether these LEA proteins contribute to the enhanced salinity tolerance of *hda19* mutants, we generated higher-order mutants. The quadruple *lea7-1/lea29-1/rab18-1/Rfhda19* mutants showed markedly reduced tolerance to 125 mM NaCl, with a survival rate of 36.7% (Fig. 5*B*), compared with 73.3% in *Rfhda19* mutants (Fig. 5*B, SI Appendix*, Fig. S2*C*). To determine whether the mRNA expression of LEA7, LEA29, and RAB18 is regulated by H3.3K27/K36ac, we analyzed mRNA levels in plants expressing HTR5 K27Q/K36Q:FLAG. Their transcript abundance was significantly elevated in these plants relative to WT, *htr4/5* mutant backgrounds, as well as in plants expressing HTR5:FLAG (Fig. 5*C*). Chromatin immunoprecipitation (ChIP)-qPCR detected the localization of both HTR5:FLAG and HTR5 K27Q/K36Q:FLAG at *LEA7, LEA29*, and *RAB18* genes (Fig. 5*D* and *E*). Together, these results indicate that H3.3 is enriched at different *LEA* genes but that only H3.3. di-acetylation at K27 and K36 results in induced LEA gene expression. The analysis of the quadruple mutant revealed that LEA7, LEA29, and RAB18 proteins promote salinity tolerance in *hda19* mutants (Fig. 6).

**Figure 5.**
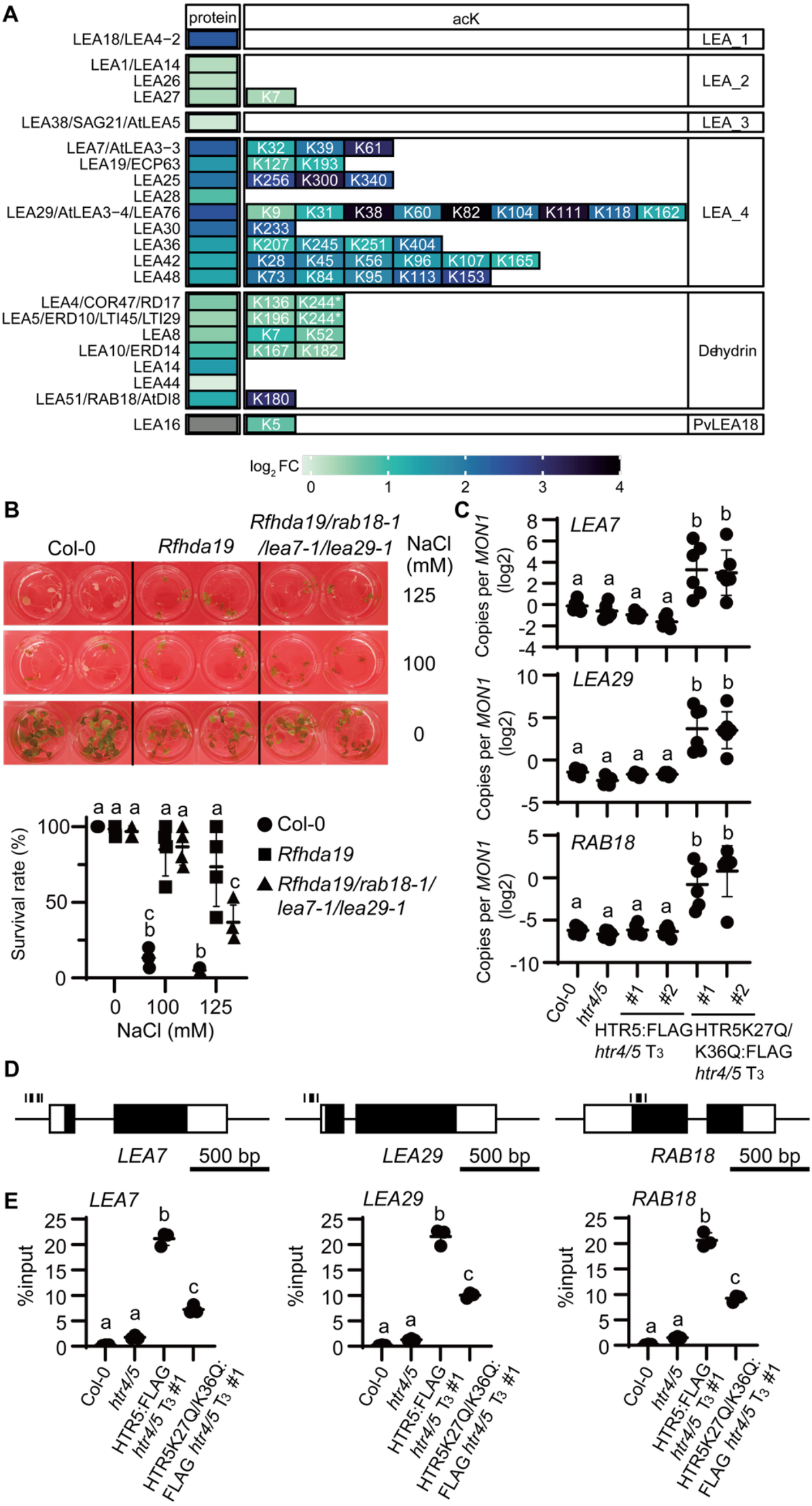
Identification of essential LEA proteins regulated by hyperacetylation of H3.3 for enhancing salinity tolerance in *hda19* mutants. (*A*) Protein abundance and lysine acetylation site (acK) levels of LEA proteins, quantified in at least 6 out of 8 replicates, are shown. The color indicates the log_2_ fold change (FC) of *hda19* versus WT Col-0. Stars next to the K position indicate sites that could belong to both LEA variants. (*B*) Deficiencies in three *LEA* genes (*RAB18*/*LEA7*/*LEA29*) decreased tolerance to salinity stress in *Rfhda19* mutants under high salinity stress conditions (125 mM NaCl). Statistical analysis was performed using a one-way ANOVA, with *P* < 0.05 considered significant (n=4). (*C*) qPCR analysis of the mRNA expression of *LEA* genes in FLAG-tagged fused HTR5 and HTR5K27Q/K36Q acetylation mimicking plants. The dataset was normalized using the reference gene (*MON1*). Statistical analysis was performed using one-way ANOVA, with *P* < 0.05 considered significant (n=6). Different letters indicate significant differences. (*D*) Schematic diagrams of primers for ChIP-qPCR of *LEA7, LEA29*, and *RAB18* genes. White boxes, black boxes, and dotted lines represent untranslated regions, exons, and amplified regions by each primer pair, respectively. (*E*) ChIP assay for FLAG-tagged fused HTR5 and HTR5K27Q/K36Q proteins using anti-FLAG antibody at the *LEA7, LEA29*, and *RAB18* genes. Statistical analysis was performed using one-way ANOVA, with *P* < 0.05 considered significant (n=3). Different letters indicate significant differences.

**Figure 6.**
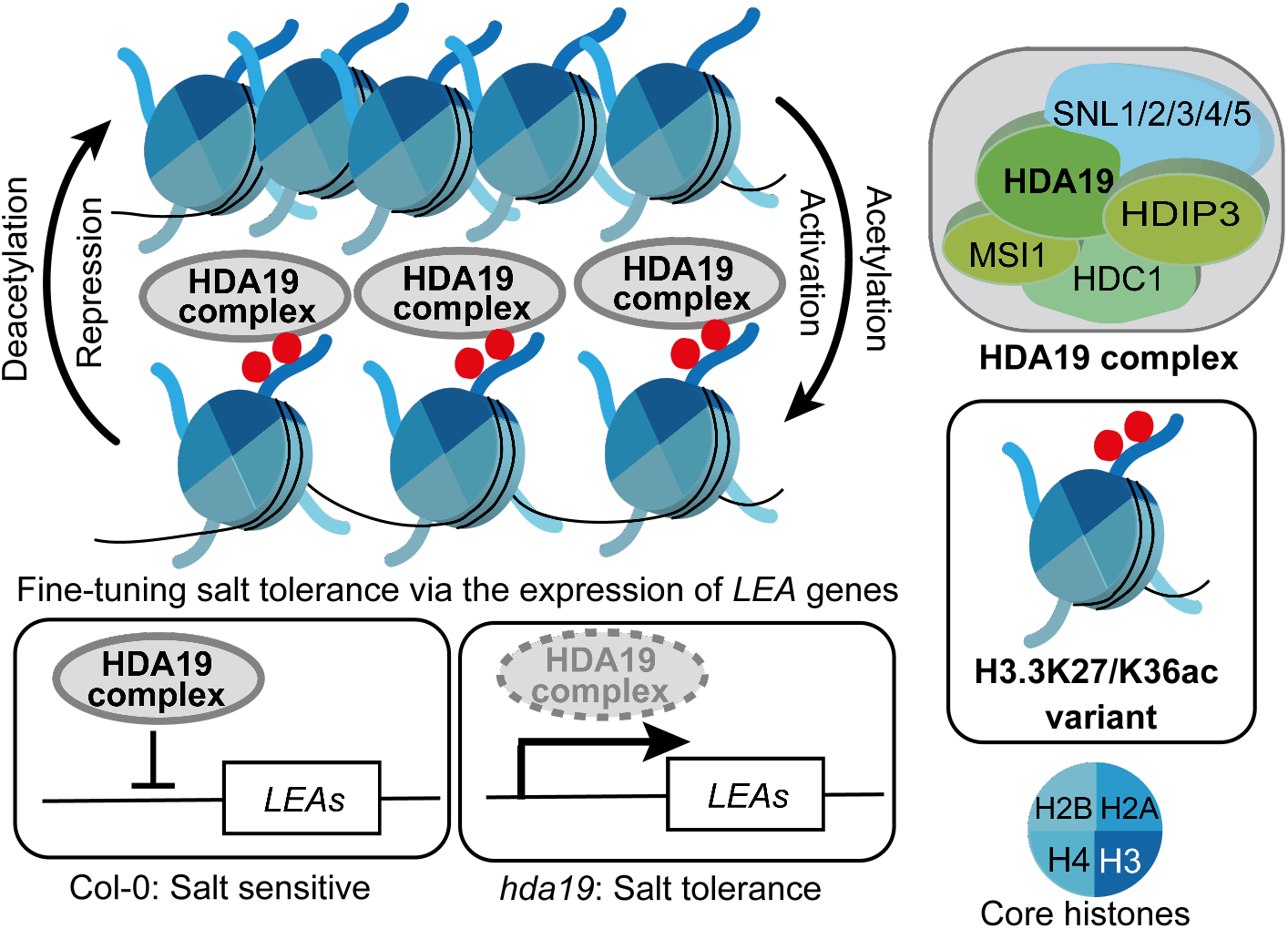
Schematic representation of a proposed mechanism of the response to salinity stress regulated by HDA19. HDA19 complex regulates the expression of *LEA* genes (e. g. *LEA7/LEA29/RAB18*) through di-acetylation of K27/K36 on histone H3.3.

## Discussion

Our study revealed that the di-acetylation site H3.3K27/K36ac is a critical epigenetic mark regulated by HDA19, which plays a key role in salinity stress tolerance in Arabidopsis. Hyperacetylation of these amino acids was found to drive the up-regulation of LEA proteins at both the transcript and protein levels in *hda19* mutants, thereby conferring salt stress tolerance.

Histone modifications represent a “histone code” that recruits transcriptional regulators to activate or repress mRNA synthesis. These histone codes are regulated by writers (catalyzing modifications), erasers (removing modifications), and reader (recognizing modifications) proteins (39, 40). Elucidating the interactors of each histone code is essential for understanding their role in the respective biological processes. The acetyltransferase GCN5 may play an antagonistic role to HDA19 at H3.3K27/K36ac sites, as it has been shown to acetylate H3K27 and K36 in Arabidopsis to activate transcription in euchromatin (41-43). The opposing photomorphogenesis phenotypes observed in *gcn5* and *hda19* mutants were absent in a double mutant (43). Furthermore, HDA19 may influence the activity of the SAGA complex, as evidenced by the hyperacetylation of its subunits TAF5 and SCS2B in *hda19* mutants. In addition to HDA19, the Arabidopsis genome encodes for three other class I HDACs (HDA6, HDA7, and HDA9) (13), although HDA7 is considered to be a pseudogene due to the of loss of the Zn^2+^ binding site in the catalytic pocket (44). HDA9, another class I HDAC, is also thought to mediate H3K27 deacetylation (45). The mono-deacetylation of H3K27ac seems to be redundantly regulated by multiple enzymes. In contrast to *hda19, hdc1* mutants are sensitive towards salt stress in comparison to WT plants (46). In *hdc1* H3K9K14ac correlates with the expression of LEA14 and RAB18 under salt stress. However, a *lea14* mutant showed an increased but not a decreased tolerance towards salt stress (46), and thus cannot be responsible for the salt-tolerance observed in the *hda19* mutants. Hence, the function of *hda19* in the salt response is different to that of *hdc1*, and further characterization of novel HDA19 substrates and interacting proteins will enhance our understanding of how plants regulate stress responses through H3.3K27/K36ac di-acetylation.

In addition to H3.3K27/K36, we identified further 48 putative substrate proteins with a consensus acetylation motif for HDA19, including 38 proteins that are not known to localize to the nucleus (Figs. 1 and 2). This raises questions about the subcellular localization of HDA19 and its substrates that require further investigation. Previously, HDA19 was reported to be exclusively localized to the nucleus (47); however maize RPD3-type HDACs have also been reported to have both a nuclear and cytoplasmic localization (48), and a significant proportion of Arabidopsis HDA9 was also found in the cytoplasm (49). Furthermore, human class I HDAC3 has been shown to reside in both the nucleus and cytosol, owing to its possession of a nuclear export signal (50). Additionally, human HDAC1-3 have been detected in the endoplasmic reticulum, where they play a role in modulating the unfolded protein response (51).

We demonstrate that three LEA proteins, which are ABA-responsive genes (36), are essential for the salt tolerance conferred by HDA19 deficiency (Fig. 5,*B*). The mapping of ChIP-seq data deposited by Liu et al (21), shows that HDA19 localizes at *LEA29* and *RAB18* genes (*SI Appendix*, Fig. S9). Combined with the ChIP-qPCR analysis of H3.3K27Q/K36Q mutants performed in this study (Fig. 5,*E*), our results suggest that HDA19 directly controls the mRNA abundance of *LEA* genes through the acetylation status of histone H3.3 K27/K36 in response to salinity stress (Fig. 6).

LEA proteins were described to accumulate during seed maturation, where they help protect cells during seed desiccation (52, 53). Expression of LEA proteins has been detected in non-plant taxa, various plant organs, and under different stress conditions, suggesting a broader role for LEA proteins in stress responses (36, 54, 55). The strong hyperacetylation of LEA proteins, particularly LEA29, implies a more direct role for HDA19 in regulation LEA proteins. Although no enzymatic functions have been reported for LEA proteins to date, multiple models have been proposed to explain their involvement in stress tolerance. These models include binding and potentially stabilizing membranes or binding directly to proteins to prevent aggregation in a chaperone-like manner (56, 57). Additionally, a molecular shield function has been suggested, where LEA proteins contribute to molecular crowding, preventing molecule collisions (58). LEA proteins, such as LEA7 and LEA29, are highly disordered in solution but form a high number of α-helices during drying (59, 60). LEA29 has been shown to form homodimers and heterodimers with LEA7 in tobacco leaves (61). The secondary structure formation of LEA7 appears to be partly dependent on the presence of membranes, although a stabilizing effect on liposomes during drying was not observed (60, 62). A protective effect of LEA7 on the activity of lactate dehydrogenase during freezing and drying was observed (62). Therefore, the accumulation and acetylation of LEA29, LEA7, and RAB18 may contribute to the protection of membranes and proteins under stress conditions. The involvement of LEA proteins in stress responses is well established, not only in Arabidopsis but also in crops (63-65). Elucidating the mode of action of these acetylated LEA proteins in the future will provide valuable insights into tolerance of salinity stress in plants.

Overall, our findings suggest that ABA signaling is the key pathway controlled by HDA19 via the regulation of histone H3.3 K27/K36 di-acetylation in response to salinity stress. Recently, angiosperm-specific HDA19-interacting proteins (HDPIs) have been identified (22), and were also found in our HDA19 seedling interactome (Fig. 4), along with other new interacting proteins. Although HDACs are highly conserved among eukaryotes, the study of plant-specific HDA19-interacting proteins, such as HDPIs, will provide further insights into the molecular mechanisms of ABA signaling regulation through H3.3 K27/K36ac.

In summary, our study identifies the histone H3.3 di-acetylation site K27/K36 as a key mediator of salinity stress tolerance, regulated by HDA19. Further characterization and identification of the coordinators of H3.3 K27/K36ac will reveal specific targets for manipulating stress responses, as HDACs represent a promising target for chemical intervention in various biological processes.

## Materials and Methods

Detailed description of plant growth conditions, the preparation of transformants, immunoblotting, RT–qPCR, ChIP-qPCR, and methods used for data analysis can be found at *SI Appendix, Materials and Methods*.

### Plant material

*Arabidopsis thaliana* (L.) Heynh. (Columbia: Col-0 as wild-type accession in this study), *hda19-3* (66); SALK_139445, *htr4htr5-1* (67); SALK_042781, *htr8-3* (67); SALK_087850, *lea7-1* (68); SALK_205942 (Col-0), *hda19-7* [the mutant has an additional base 79G inserted in the *HDA19* gene and is a novel allele is resulting from a progeny of genome-edited Col-0 plants using the same guide RNA as reported in our previous study (16)], Reverted fertility in *hda19-7* (*Rfhda19*) (see for details *SI Appendix*, Fig. S1*A*), *lea29-1* (in 104T, a genome-edited allele) (*SI Appendix*, Fig. S10*A*) and *rab18-1* (in 223A, a genome-edited allele) (*SI Appendix*, Fig. S10*B*) and were used in the course of this study.

### Evaluation of salinity tolerance

When plants were 5 days old (counted from seed germination), 20 µL or 25 µL of 5 M NaCl were added to each well, yielding final concentrations of 100 mM or 125 mM NaCl per well, respectively, while control plants received no salt. Percent survival was assessed five days later using three biological replicates, each containing 15 plants. Plants with green true leaves were scored as survivors. Survival rates were statistically analyzed by a one-way ANOVA in the GraphPad Prism version 10. 2. 3 for Windows (GraphPad Software).

### Primary antibodies for Immunoblotting and ChIP-qPCR

Primary antibody dilutions were as follows: H3.3 (HTR4/HTR5/HTR8) K27/K36ac, 1: 500 (polyclonal antibody raised against H3.3K27acK36ac in this study); H3, 1:2,000 (polyclonal, Abcam, Cambridge, MA, 1791); and Myc-tag, 1:1,000 (monoclonal, MBL, Japan M185-3L). Synthetic peptides [CKAARKAcSAPTTGGVKAcKP] were used as antigen to produce polyclonal antibodies against H3.3K27/K36ac (Scrum, Tokyo, Japan).

### Enrichment of active HDACs using AsuHd probes

Plant material was homogenized and suspended in extraction buffer (EB) containing 50 mM HEPES (pH 7.5), 150 mM NaCl, 10% [v/v] glycerol, 5 mM ascorbate, 0.25% [v/v] Triton X-100, and cOmplete™ EDTA-free Protease Inhibitor Cocktail (Roche). Extracts were incubated at 4 °C for 1 h with constant agitation, and cell debris was pelleted by centrifugation at 18,000 x *g* for 25 min at 4 °C. Protein concentrations in the supernatants were determined using the Pierce 660 nm Protein Assay (Thermo Fisher Scientific). Resins with mini-AsuHd (HDAC trap) and mini-Lys peptide (control) probes (23), were equilibrated 3x with 600 µL wash buffer (WB) (50 mM Hepes pH 7.5, 150 mM NaCl, 10% [v/v] glycerol, 5 mM ascorbate). Equal amounts of each sample (2.5 mg) were incubated overnight at 4 °C with 10 µL of both mini-AsuHD and mini-Lys beads under constant agitation. Beads were transferred onto a micro-centrifugal filter system (amchro GmbH) and washed 5x with 250 µL WB. An on-bead trypsin digestion was performed as described previously (28). Peptides were desalted using C18 Stage tips (69) prior LC-MS/MS analysis.

### Myc pulldown

Plant material was homogenized and suspended in EB buffer as described above. EZview™ Red Anti-c-Myc Affinity Gel (Merck Millipore, Darmstadt, Germany) was equilibrated 3x with WB (50 mM Tris pH 7.5, 150 mM NaCl, 10% [v/v] glycerol, 2 mM EDTA). Protein extracts were incubated on beads over-night at 4 °C with constant agitation. Beads were collected by slow centrifugation (500 x *g*) at 4 °C and washed 3x with 1 mL WB for 5 min with constant agitation. Proteins were eluted two times with 30 µL 0.1% (v/v) TFA for 5 min and collected by gentle centrifugation. An in-solution trypsin digestion was performed as described previously. For total proteome analysis, peptides from input samples were prepared using small-scale filter-aided sample preparation (FASP) as described previously (70).

### Quantitative proteomics and lysine acetylome

Proteins were extracted from plant material under denaturing conditions as described previously (71). DTT was replaced with tris(2-carboxyethyl)phosphine (TCEP). Proteins were processed using single-pot, solid-phase-enhanced sample preparation (SP3) (72). After the tryptic digest, peptides were labeled using tandem mass tags (TMT) (73). TMT10plex™ (Thermo Fisher Scientific) reagents were used. Samples were randomized within each TMT set. Each channel was assigned to one genotype or treatment, or a pool of all samples for normalization purposes. Peptides were resuspended in TBS buffer (50 mM Tris–HCl, 150 mM NaCl, pH 7.6). A more than 98% labeling efficiency was confirmed by LC-MS/MS analysis and samples were pooled afterwards. 30 µg of pooled peptides was stored for whole proteome analysis. Lysine-acetylated peptides were enriched using PTMScan® Acetyl-Lysine Motif [Ac-K] Immunoaffinity Beads (Cell Signaling Technology) as described previously (71). All total proteome and acetylome samples were desalted using SDB-RPS Stage tips (71), eluted in three separate fractions, and dried using a vacuum concentrator.

### LC-MS/MS data acquisition

Peptides were resuspended in 2% (v/v) acetonitrile and 0.1% (v/v) TFA. Aceylome, proteome and HDAC pulldown samples were measured using an EASY-nLC 1200 system coupled to an Exploris 480 mass spectrometer (Thermo Fisher Scientific). Myc pulldown and corresponding input samples were measured using an EASY-nLC 1200 system coupled to an Q Exactive HF mass spectrometer (Thermo Fisher Scientific). The peptides were separated on 20 cm fritless silica emitters with a 0.75 µm inner diameter (CoAnn). Emitters were packed in-house with reversed-phase 1.9 μm ReproSil-Pur C_18_-AQ (Dr Maisch GmbH). A 115 min (78 min for Myc pulldown) LC gradient using 0.1% (v/v) formic acid as buffer A and 80% (v/v) acetonitrile, 0.1% (v/v) formic acid as buffer B was used. Data acquisition settings for all experiments are listed in *SI Appendix*, Table S7.

## Acknowledgments

The authors are grateful to Ms. S. Nakae, Ms. K. Kaneko, and Ms. F. Sakai in the support unit for Bio-Material Analysis and RIKEN CBS Research Resources Division for technical help with the nucleotide sequencing analyses. We also thank Prof. M. Matsui and Dr. W.D. Ong at RIKEN for providing the *lea7-1* as well as Ms. A. Sato (Plant Genomic Network Research Team), Ms. T. Kaizuka (Drug Discovery Seeds Development Unit), Dr. K. Matsumoto (Drug Discovery Seeds Development Unit), Dr. A. Idei (Drug Discovery Seeds Development Unit) for technical support. We would like to thank Mrs. Paulina Heinkow for technical support with the LC-MS/MS analysis within the MSPUB proteomics unit of the University of Muenster. This work was supported by the following grants and funding agencies: ASPIRE JPMJAP24A3 (MS), CREST JPMJCR13B4 (MS), GteX programs of the Japan Science and Technology Agency JPMJGX23B0 (MS), “Epigenome Manipulation Project” and “Global Commons Project” of RIKEN (MS), the Ministry of Education Culture, Sports and Technology of Japan KAKENHI Grant 19K05960 (MU), and the Deutsche Forschungsgemeinschaft (DFG, German Research Foundation) – INST 211/744-1 FUGG for instrumentation (IF).

## Author Contributions

M.U., M.S., D.S., and I.F. designed research; F.K., M.U., A.I., S. T., K.S., H.T., S.T., J.E., J.I., J.S., and T.A. performed research; F.K. and M.U. analyzed data; and F.K., M.U., M.S., and I.F. wrote the paper.

## Data and materials availability

Mass spectrometry proteomics data have been deposited at the ProteomeXchange Consortium (https://proteomecentral.proteomexchange.org/ui) via the JPOSTDB partner repository (https://repository.jpostdb.org/) with the identifiers JPST004010 and PXD067353.

Gene identifiers and sequence information can be found in The Arabidopsis Information Resource-TAIR database (www.arabidopsis.org) under the following accession numbers: HDA19 (AT4G38130), HTR4 (AT4G40030), HTR5 (AT4G40040), HTR8 (AT5G10980), LEA7 (AT1G52690), LEA29 (AT3G15670), RAB18 (AT5G66400). Plasmids and seeds are available from M.S. upon reasonable request.

