## Supplementary material for "HDA19-mediated deacetylation of histone H3.3 lysine 27 and 36 regulates plant sensitivity to salt stress": SI Appendix

**This PDF file includes:**

Supporting text
Figures S1 to S11
Tables S1 to S7
Legends for Datasets S1 to S5
SI References

**Other supporting materials for this manuscript include the following:**
Datasets S1 to S5

**Supporting Information Text**

Materials and methods for plant growth conditions, the preparation of transformants, immunoblotting, chromatin immunoprecipitated (ChIP)-quantitative polymerase chain reaction (qPCR), Reverse transcription quantitative polymerase chain reaction (RT-qPCR), and methods used for data analysis.

**Plant growth conditions.** Plants treated with trichostatin A (T1952, Sigma–Aldrich, St. Louis, MO) were prepared as described previously (1). After surface-sterilization with sodium hypochlorite, followed by two rinses with distilled water, seeds were floated on 1 mL liquid media (half-strength MS medium with 0.5% MES and 0.1% agar, pH 5.7) at 4 °C for 48 h in 24-well (Inside diameter: 15.4 mm) tissue culture plates (TPP; Trasadingen, Switzerland). After germination, the plants in the 24-well tissue culture plates were placed in a growth chamber at 22 °C with a long-day photoperiod (16h/8h light/dark cycle) at 50–100 μE $m^{-2}$ $s^{-1}$. For the acetylome and pull-down analyses, 5-d-old-plants were harvested, and for the proteome analysis of the early salt stress response, plants were exposed to salinity stress for 2 h before harvesting. After harvesting of four biological replicates each, plants were immediately flash frozen in liquid nitrogen and stored at -80 °C. The plant samples for proteomic analysis were dried using a freeze dryer (FRD-830D, IWAKI CO., LTD., Tokyo, Japan) for 24 hours. Developmental stages of stamen and pistil in *Rfhda19* and wild-type plants were referred to Smyth et al 1990 (2).

**Generation of mutant lines using the CRISPR/Cas9 system.** The *rab18-1* and *lea29-1* alleles were generated by genome editing as described previously (*SI Appendix*, Fig. S9)(3, 4). The sgRNAs used for mutagenesis were designed using the CRISPRdirect program (5). To express a single guide RNA (sgRNA) and CRISPR/Cas9 protein, pZH_OsU3gYSA_FFCas9 and pUC_AtU6oligo vectors were used for targeted mutagenesis in *RAB18* and *LEA29*. The sgRNA information used to target them is presented in *SI Appendix*, Table S4.

**Generation of complement lines expressing H3.3 K/Q mutant.** K to Q substitutions in K27 and K36 of HTR5 coding sequence were introduced by PCR-based site-directed mutagenesis (6), and fused in reading frame with a C-terminal FLAG (DYKDHDGDYKDHIDYKDDDDK). Genomic DNA of HTR5 was used and the construct was first cloned into a pENTR/DTOPO vector (Thermo Fisher Scientific, Plaquemine, LA). The resultant DNA was used as a template for mutagenesis. The HTR5K27Q/K36Q:FLAG coding gene was expressed under the control of its own promoter and terminator in the *htr4htr5-1* mutant background as follows. Amplified PCR products of promoter, gene body, and terminator fragments were assembled into plant expression vector pHGW (7), by HiFi DNA Assembly (New England Biolabs, Ipswich, MA). Primer pairs used for mutagenesis and cloning are shown in *SI Appendix*, Table S4. Agrobacterium tumefaciens strain C58C1 was transformed by the pHGW binary vector and used for floral dip transformation in *htr4/5* according to the method of Clough and Bent (8).

**Immunoblotting.** Tissue powder from ten five-day-old plants was solubilized in 2 x Laemmli buffer (100 μl/ 10 plants) under reducing conditions (200 mM DTT or 2% β-mercaptoethanol) and heated at 95 °C for 3 min. Protein extracts were separated by sodium dodecyl sulfate polyacrylamide gel electrophoresis on 15% bis-Tris gels (Nacalai Tesque) and subsequently transferred to TransBlot Turbo mini-size PVDF membranes (Bio-Rad, Hercules, CA). The membranes were blocked for 1 h with 5% skim milk, incubated overnight at 4 °C with the primary antibody, and then incubated for 1 h with the anti-rabbit secondary antibody (NA934, Cytiva) or anti-mouse (NA931, Cytiva) IgG secondary antibody conjugated to horseradish peroxidase. Immunoreactive proteins were detected using Chemi-Lumi One Super (Nacalai Tesque, Japan), and images were acquired with an ImageQuant800 scanner (Cytiva). Band intensities were quantified with ImageJ software. Equal protein loading was verified by probing for histone H3. To visualize the levels of acH3.3K27/K36 in *Rfhda19* and *hda19-3* in Fig. 3C, fluorescence immunoblotting was conducted using low fluorescence PVDF (Bio-Rad) and IRDye 800CW donkey anti-rabbit IgG secondary antibody (LI-COR Biosciences), according to the manufacturer's instructions. Images were acquired with an Odyssey imager and Image Studio software (LI-COR Biosciences).

**RT-qPCR analysis.** First-strand cDNA was synthesized from 250 ng total RNA extracted from 10 plants per condition using random primers and ReverTra Ace (TOYOBO). Transcript levels were analyzed using Fast SYBR Green Master Mix (Applied Biosystems, Foster City, CA) and the StepOnePlus Real-Time PCR System (Applied Biosystems) according to the manufacturer's protocols and previous study (9). Gene-specific primers were designed using the PrimerQuest tool (http://sg.idtdna.com/primerquest/Home/Index). The relevant primers are listed in *SI Appendix*, Table S5. Changes in gene expression (n=6) were statistically analyzed by a one-way ANOVA in the GraphPad Prism version 10. 2. 3 for Windows (GraphPad Software).

**ChIP-qPCR**. ChIP assay was performed according to the method for the *A. thaliana* as previously described (10). Covaris S220 ultrasonicator was used for sonication with Covaris microTUBEs (Covaris) (fragmentation condition: 16 min, Peak power: 140, Duty Factor: 5, Cycle Burst: 200). 200-300mg five-day-old seedlings were used for the ChIP assay. Anti-FLAG mono-clonal antibody (M185-3L, MBL, Japan) was used in this study. Three technical repeats were performed. The precipitates were analyzed with qPCR. The primers used are listed in *SI Appendix*, Table S6. ChIP-seq data of HDA19-FLAG available from Liu et al. (11), was referred to design the primer pair for *LEA7* and *LEA29* (*SI Appendix*, S9). The primer pair for *RAB18* was designed in Perrella et al. (12). Changes in the recovery (% of input) (n=3) were statistically analyzed by a one-way ANOVA in the GraphPad Prism version 10. 2. 3 for Windows (GraphPad Software).

**Data analysis.** LC-MS/MS raw data were analyzed with the MaxQuant (MQ) software (v 2.4.14.0, https://maxquant.net) (13). The raw data was searched against an in house curated Araport11 database including organellar specific proteins after processing based on targetP2 (14), and a database containing mitochondria-encoded proteins after RNA editing (15, 16). Common contaminants lists and a reverse decoy database were enabled in the MaxQuant software settings. Proteome and acetylome samples of *hda19-3* and *Rfhda19* were analyzed in the same MQ search using dedicated parameter groups. All pull-down samples and respective inputs were analyzed in. the same MQ search using dedicated parameter groups. Carbamidomethylation of cysteines was set as fixed modification and oxidation of methionine and acetylation of protein N-termini were set as variable modifications. Trypsin was selected as protease. For acetylome samples, lysine acetylation was set as variable modifications, the number of max. missed cleavages was set to 3 (2 for all other samples), and PTM was set as TRUE. Match between runs was activated for all samples. min. score for modified peptides was set to 35 for the acetylome search as reported previously (17). $MS^2$-based quantification was activated and set to TMT10plex for acetylome and corresponding proteome samples. TMT correction factors were adjusted to the specific TMT batch, filter by PIF was activated, and normalization was set to "Weighted ratio to reference channel". Re-quantify was enabled, as well as separate variable modifications for first search. For pull-down and respective input samples label-free quantification (LFQ) was enabled and the LFQ min. ratio count was set as 1. Separate LFQ in parameter groups was activated. iBAQ was enabled. min. peptide length was set to 7 for all searches. Both PSM and protein FDR were set to 0.01.
MQ output tables were imported into Perseus (v 2.0.11.0, https://maxquant.net/perseus/) (18). Contaminants and reverse hits were removed by filtering. Proteins only identified by site were removed by filtering for pull-down samples only. LFQ intensities and corrected reporter intensities were $\log_2$ transformed. Protein groups and acK sites were filtered for minimum valid values as indicated in DataS1-5. TMT samples were normalized by median subtraction. Missing values were imputed following normal distribution in Myc input samples (width = 0.3, down shift = 1.8), Myc pull-down WT control (width = 0.5, down shift = 2.2), and HDAC pull-down mini-Lys controls (width = 0.4, down shift = 1.5). Data was exported from Perseus and imported into R (v 4.4.1). Statistical analysis was performed using the *limma* package (v 3.60.6) (19). The *p*-value was adjusted with Benjamini-Hochberg. Packages dplyr (v 1.1.4), tidyr (v 1.3.1), reshape2 (v 1.4.4), stringi (v 1.8.4), and stringr (v 1.5.1) were used for data processing. Results were visualized using the packages ggplot2 (v 3.5.2) and ggrepel (v 0.9.6). SUBAcon annotations of protein subcellular localizations were retrieved from the SUBA5 database (20). Sequence logo of significantly upregulated acK sites (*hda19* vs. WT, $\log_2$ FC > 1, FDR < 0.05) was analyzed using the iceLogo software (21). The unregulated acK sites with a $\log_2$ FC < 1 were used as a negative set. A *p*-value cut-off of 0.05 was used. Functional enrichment analysis was performed using the Proteins with Values/Ranks function of stringDB (22). A rank value was calculated by multiplying the $\log_2$ acK FC with the $-\log_{10}$(adjusted

*p*-value). For proteins with multiple acK sites, the site with the highest rank was used. GO cellular component, GO biological process, KEGG, UniProt Keywords were considered. Only terms showing an increase in their abundance were used. String Network analysis was done using Cytoscape (v 3.10.2) together with the stringApp (v 2.2.0) and the Omics Visualizer (v 1.3.1)(23-25). The confidence cut-off was set to 0.7. Histone variants have been annotated based on Probst et al., 2020 (26), and LEA proteins based on Hundertmark and Hincha, 2008 (27). The localization of LEA proteins was referred to Candat et al 2014 (28).

**Figures**

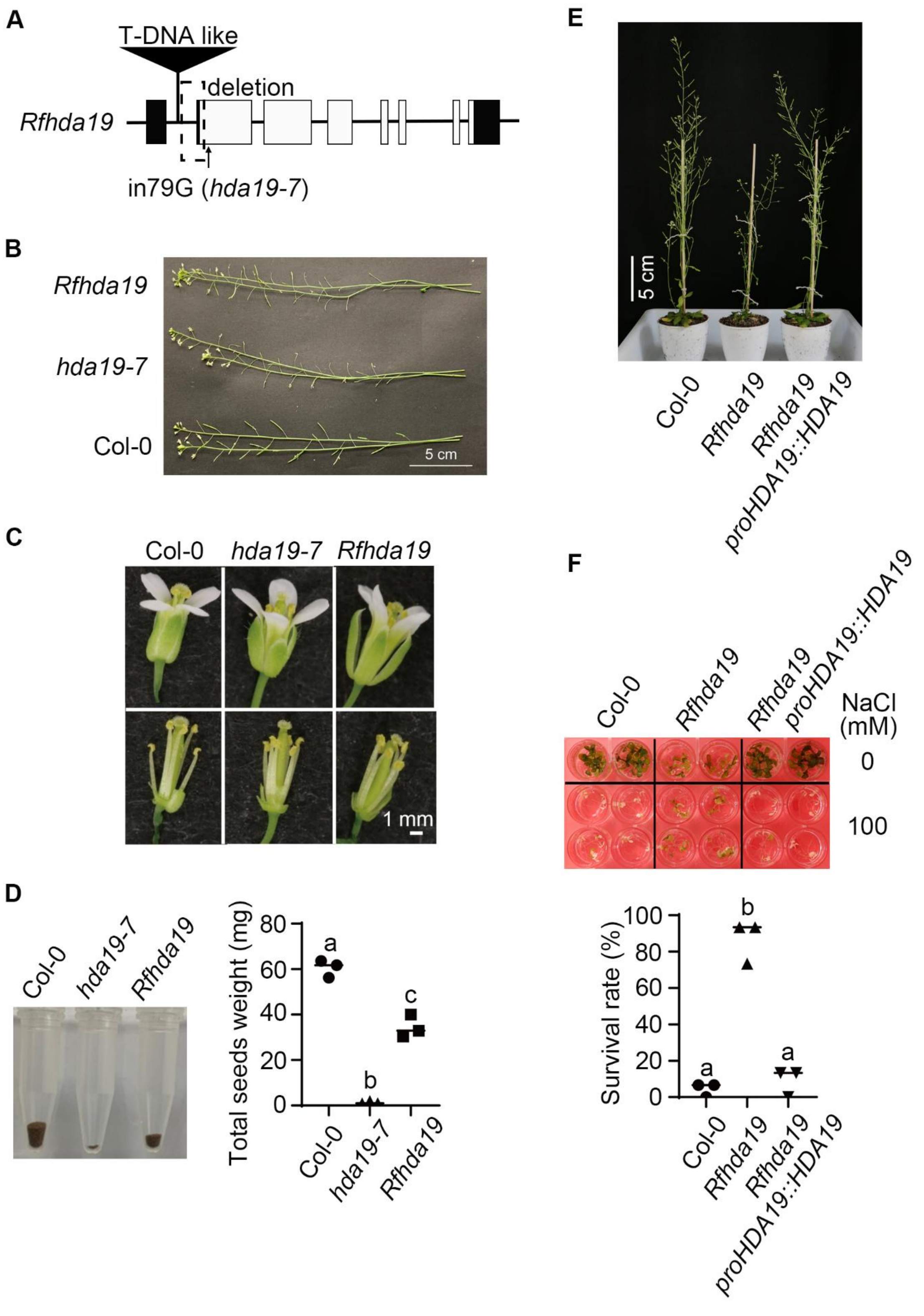


**Fig. S1.** Characterization of *Rfhda19* (Reverted fertility in *hda19*). (*A*) Genome rearrangements around the *HDA19* locus in *hda19-7*. (*B*) Morphological phenotype of the siliques of the main stem in *Rfhda19*. (*C*) Morphological phenotype of stamen and pistil in *Rfhda19* at stage14, as defined Smyth et al 1990 (2). (*D*) Comparison of seed numbers harvested from three individual plants of *Rfhda1*9*.* (*E*) Complementation of *Rfhda1*9 with a HDA19 genomic fragment. (*F*) Loss of salinity stress tolerance upon complementation of HDA19 in *Rfhda1*9. Statistical analyses in D and F were performed using a one-way ANOVA test ($P$ < 0.05, n=3). Different letters indicate significant differences.

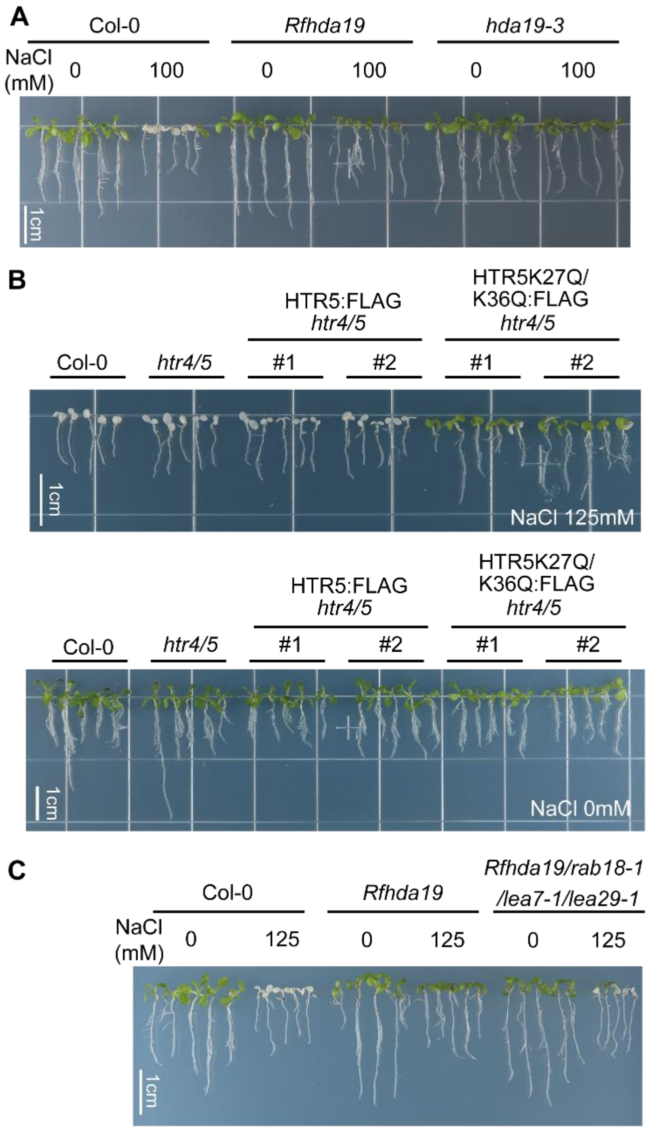


**Fig. S2. Vertical pictures of different genotypes in response to salinity stress.** (*A*) *Rfhda19* and *hda19-3* exhibited similar tolerance to salinity stress (100 mM NaCl). Corresponding to Fig. 1A. (B) Expression of FLAG-tagged HTR5K27Q/K36Q acetylation mimicking variant in *htr4/5* enhanced tolerance to salinity stress (125 mM NaCl). Corresponding to Fig. 3D. (C) Loss of three *LEA* genes (*RAB18*/*LEA7*/*LEA29*) decreased salinity stress tolerance in the *Rfhda19* mutant background under high salinity stress conditions (125 mM NaCl). Corresponding to Fig. 5B.

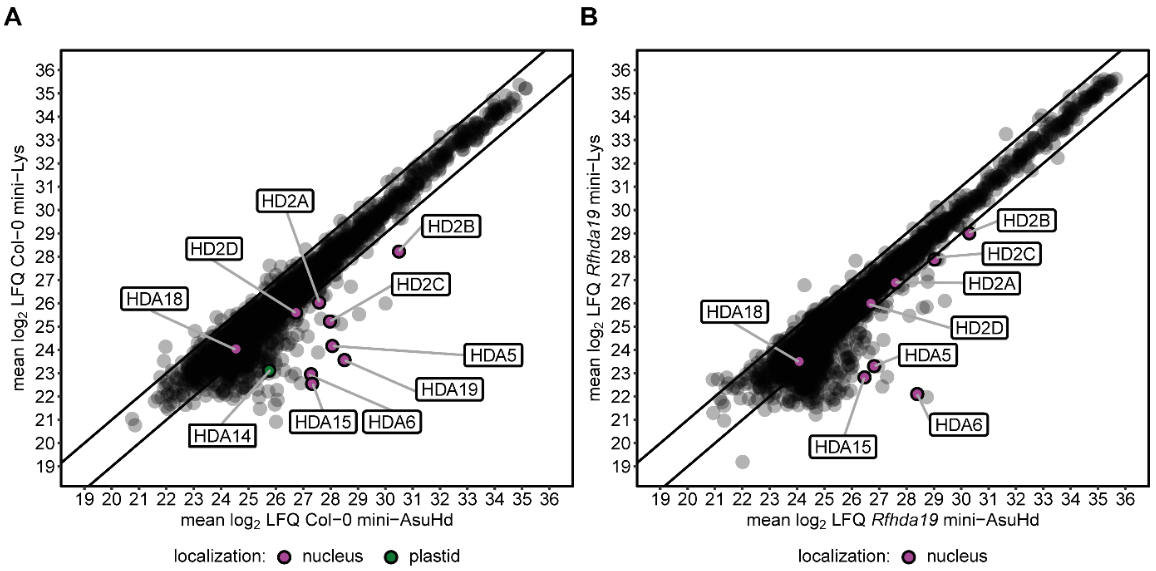


**Fig. S3.** *Rfhda19* is a lacking an active HDA19 enzyme. (*A*-*B*) Scatter plots comparing mean $\log_2$ LFQ values of mini-AsuHd and mini-Lys pull-downs in (A) Col-0 WT and (B) *Rfhda19* (n = 2). All quantified HDACs are labeled and colored according to their subcellular localization based on SUBA5 (20). The original data can be found in Dataset S1.

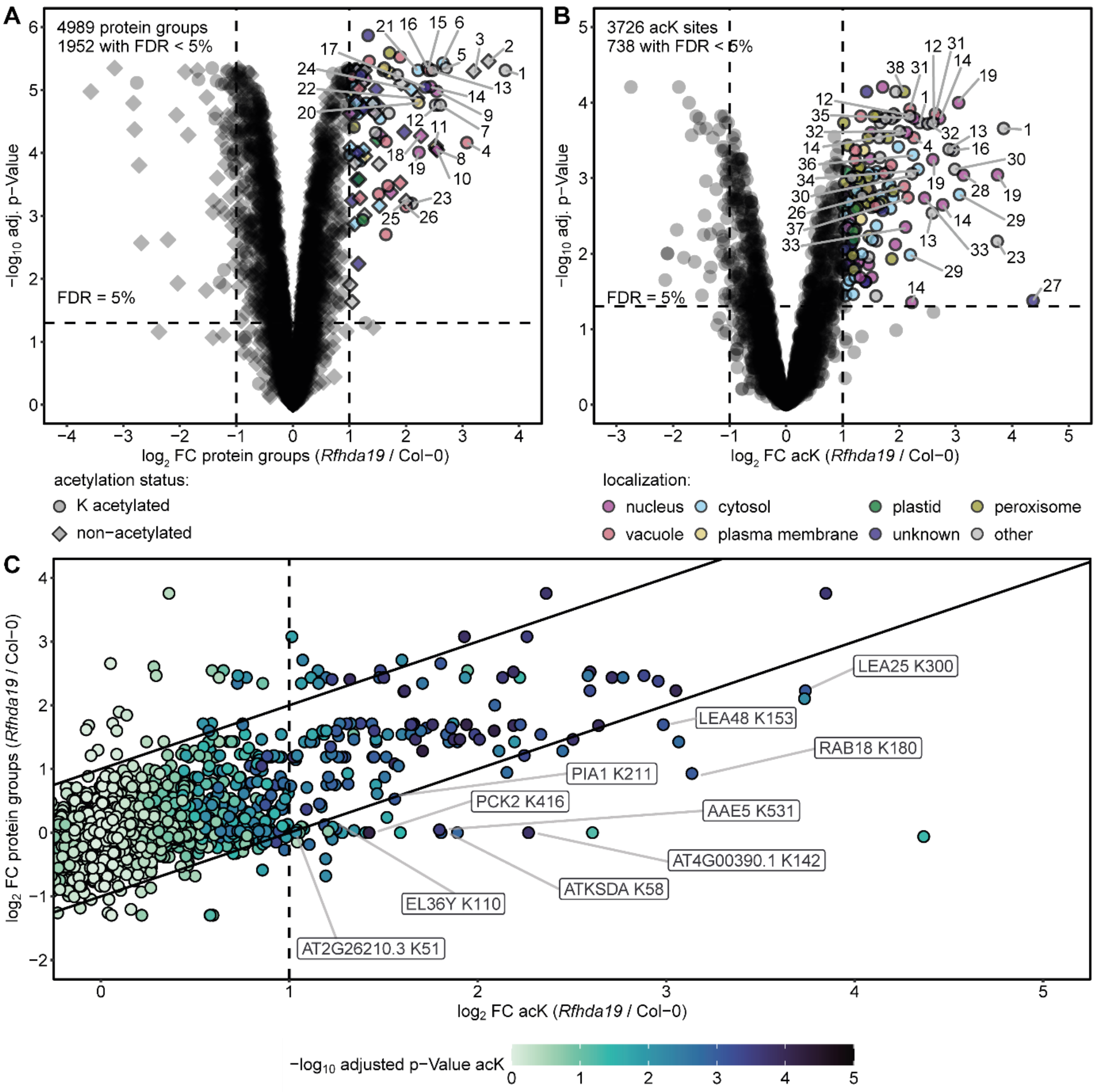


**Fig. S4.** Quantitative proteome and lysine acetylome of *Rfhda19* compared to Col-0 WT. (*A*-*B*) Volcano plots showing (*A*) protein groups and (*B*) lysine acetylation (acK) site $\log_2$ fold abundance changes in *Rfhda19* compared to Col-0 WT, along with -$\log_{10}$(adjusted p-values) from *limma* analysis (16). The p-values were adjusted using a Benjamini-Hochberg correction. Only acK sites and protein groups quantified in at least 2-4 biological replicates were included in the analysis. $\log_2$ fold changes (FC) of +/- 1 and false discovery rates (FDR) of < 5 % are indicated by dashed lines. Significantly altered proteins and acK sites ($\log_2$ FC > 1, FDR < 5 %) are colored according to their SUBA5 localization (20). AcK sites and proteins with a $\log_2$ FC > 2 are numbered. Protein numbers with symbols or identifiers from 1 to 38: 1. CRU4, 2. AT2G05580, 3. CRU2, 4. LEA18, 5. PAP85, 6. TASTY, 7. AT4G36700, 8. RD29B, 9. LEA7, 10. AT2G18540, 11. AT3G20730, 12. CRU3, 13. CRU1, 14. LEA29, 15. OELO2, 16. LEA30, 17. OELO4, 18. AT2G29065, 19. LEA25, 20. ATS3, 21. PER1, 22. AT2G28490, 23. SESA1, 24. ATXIE1, 25. AT5G35660, 26. SESA3, 27. DRP1A, 28. RAB18, 29. ECP63, 30. LEA48, 31. OLEO1, 32. LEA42, 33. SLDP1, 34. LEA36, 35. AT4G00390, 36. PCK1, 37. OLEO3, 38. mLS. (*C*) Scatter plot of $\log_2$ acK FC against $\log_2$ protein FC of *Rfhda19* vs WT. Only acK sites with a $\log_2$ FC > 0 are shown. Colors indicate the -$\log_{10}$(adjusted p-value) of acK site changes. $\log_2$ acK FC > 1 is indicated by a dashed line. Diagonal lines (slope = 1, intercepts = +/- 1) represent an absolute fold change difference of 1 between acK sites and proteins. Only proteins/acK sites which are also labeled in Fig. 2F are labeled here. The original data can be found in Dataset S2.

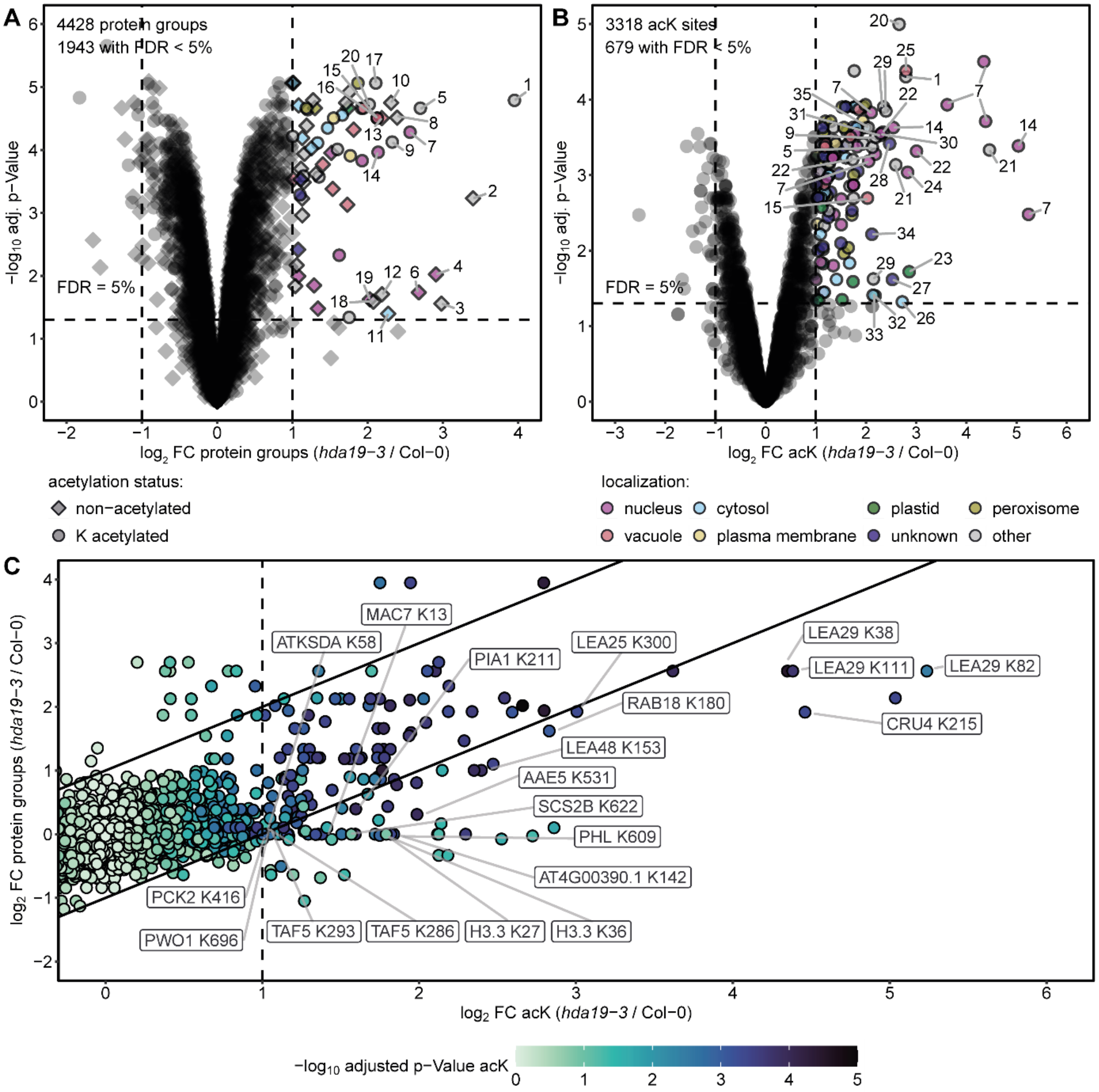


**Fig. S5.** Quantitative proteome and lysine acetylome of *hda19-3* compared to Col-0 WT. (*A*-*B*) Volcano plots showing (*A*) protein groups and (*B*) lysine acetylation (acK) site $\log_2$ fold abundance changes in *hda19-3* compared to Col-0 WT, along with $-\log_{10}$(adjusted p-values) from *limma* analysis (16). The p-values were adjusted using a Benjamini-Hochberg correction. Only acK sites and protein groups quantified in at least 2-4 biological replicates were included in the analysis. $\text{Log}_2$ fold changes (FC) of +/- 1 and false discovery rates (FDR) of < 5 % are indicated by dashed lines. Significantly altered proteins and acK sites ($\log_2$ FC > 1, FDR < 5 %) are colored according to their SUBA5 localization (20). AcK sites and proteins with a $\log_2$ FC > 2 are numbered. Protein numbers with symbols or identifiers from 1 to 35:
1. AT2G05580, 2. TRP3, 3. GRP3, 4. GRP2B, 5. CRU3, 6. GRP2, 7. LEA29, 8. PAP85, 9. CRU1, 10. AT4G36700, 11. RPS13, 12. AT1G47540, 13. CLO1, 14. LEA7, 15. OLEO2, 16. OLEO4, 17. SBTI1.1, 18. HEL, 19. PUX7, 20. LEA30, 21. CRU4, 22. LEA25, 23. NIR, 24. RAB18, 25. OLEO1, 26. AXS2, 27. LOS1, 28. ChlADR, 29. LEA48, 30. TAF14B, 31. PER1, 32. RBCS1A, 33. LOS2, 34. TAF5, 35. SESA1. (*C*) Scatter plot of log2 acK FC against log2 protein FC of *hda19-3* vs WT. Only acK sites with a log2 FC > 0 are shown. Colors indicate the -log10(adjusted p-value) of acK site changes. log2 acK FC > 1 is indicated by a dashed line. Diagonal lines (slope = 1, intercepts = +/- 1) represent an absolute fold change difference of 1 between acK sites and proteins. Only

proteins/acK sites which are also labeled in Fig. 2F are labeled here. The original data can be found in Dataset S2.

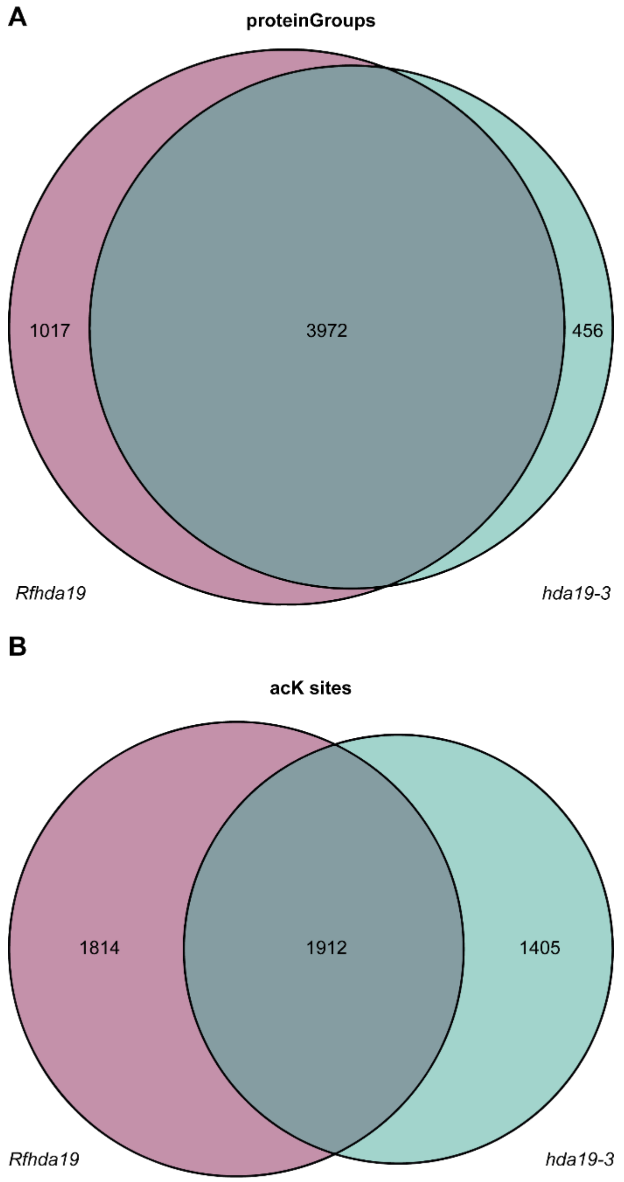


**Fig. S6.** Venn diagrams showing overlap of proteomes between *Rfhda19* and *hda19-3*. Overlap of (*A*) protein groups and (*B*) lysine acetylation (acK) sites between *Rfhda19* and *hda19-3*. Proteins and acK sites quantified in at least two out of four biological replicates were used. The original data can be found in Dataset S2 and Dataset S3.

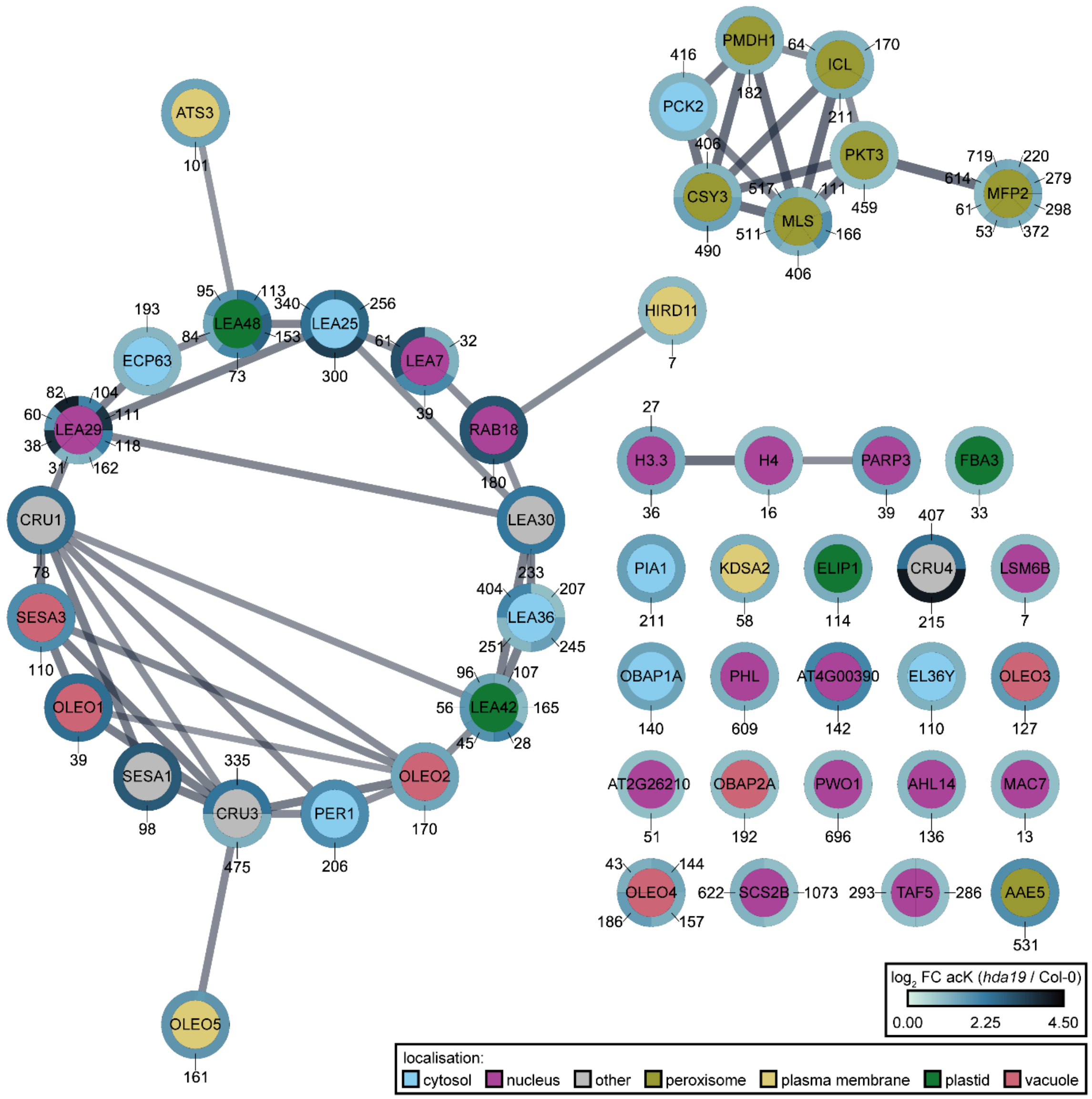


**Fig. S7.** STRING network of hyperacetylated proteins in *hda19*. The network was generated using Cytoscape (23) and includes hyperacetylated proteins ($\log_2$ FC acK > 1, FDR < 0.05) identified in the combined analysis of *hda19-3* and *Rfhda19*. The outer circle color represents $\log_2$ FC of acK sites according to the color scale, with blue indicating higher values. The position of each acK site is indicated by the outer labels. For histone proteins, lysine positions have been adjusted to reflect their position after removal of the starter methionine. The circle inner color represents subcellular localization, as annotated by SUBA5 (20). The localization of LEA proteins was updated based on Candat et al 2014 (28). For proteins with unknown localization due to splice forms not present in SUBA5, the localization of the first splice form was assumed. The original data is available in DataS4.

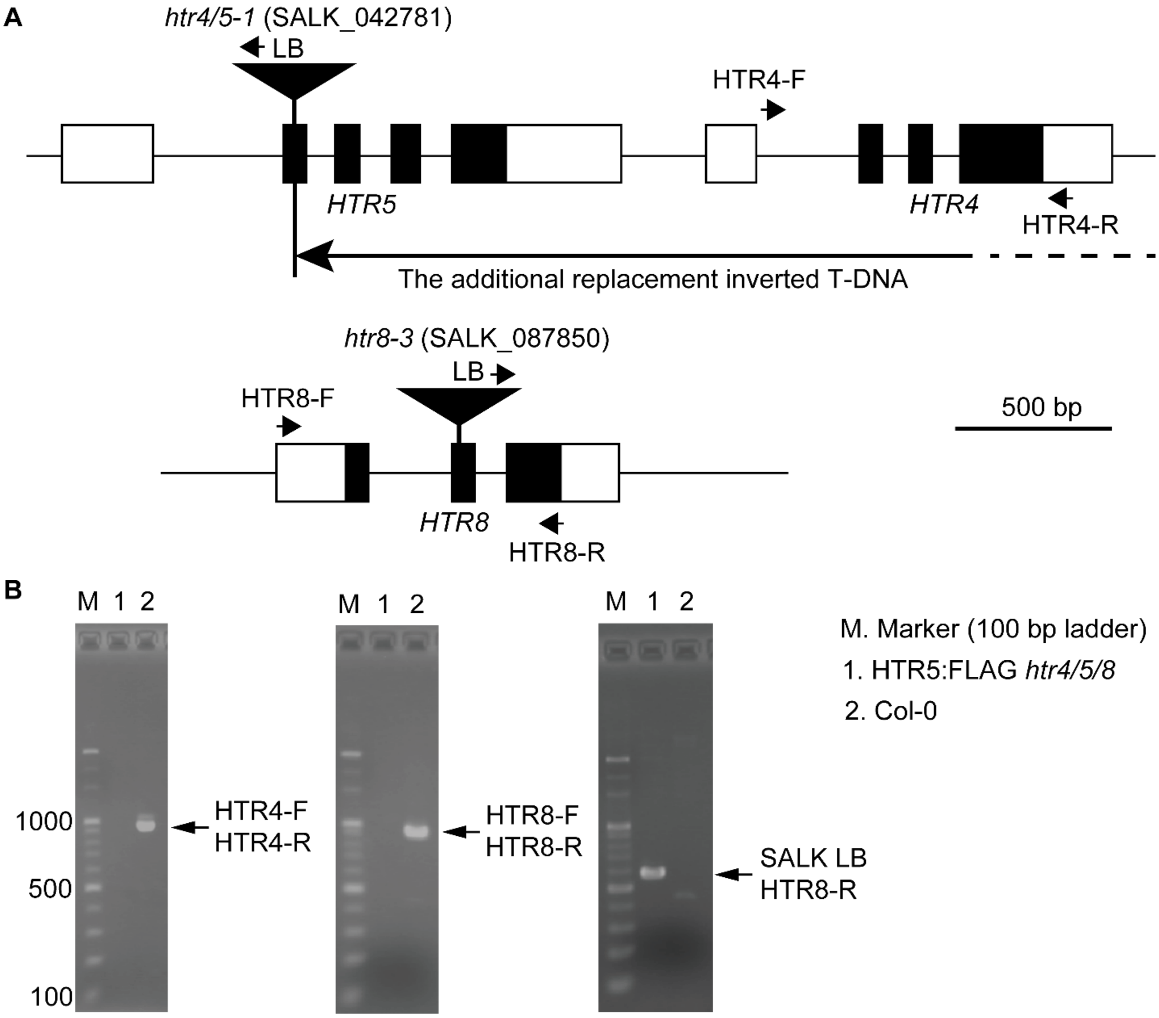


**Fig. S8.** Confirmation of *htr4/5/8* triple mutant used to generate a complement line rescued by HTR5:FLAG expression. (*A*) Schematic representation of the *HTR4*, *HTR5*, and *HTR8* gene structures and the T-DNA insertions in *htr4/5-1* (SALK_042781) and *htr8-3* (SALK_087850) according to (29). Black boxes, white boxes, and triangles represent exons, untranslated regions, and T-DNA, respectively. Arrows represent primer pairs used for PCRs in B. The information on primer sequences used for B is available from Liu et al 2024 (29). (*B*) Gel electrophoresis of PCR products for a T-DNA insertion site in *htr8-3,* and deletion of *HTR4* in *htr4/5-1*.

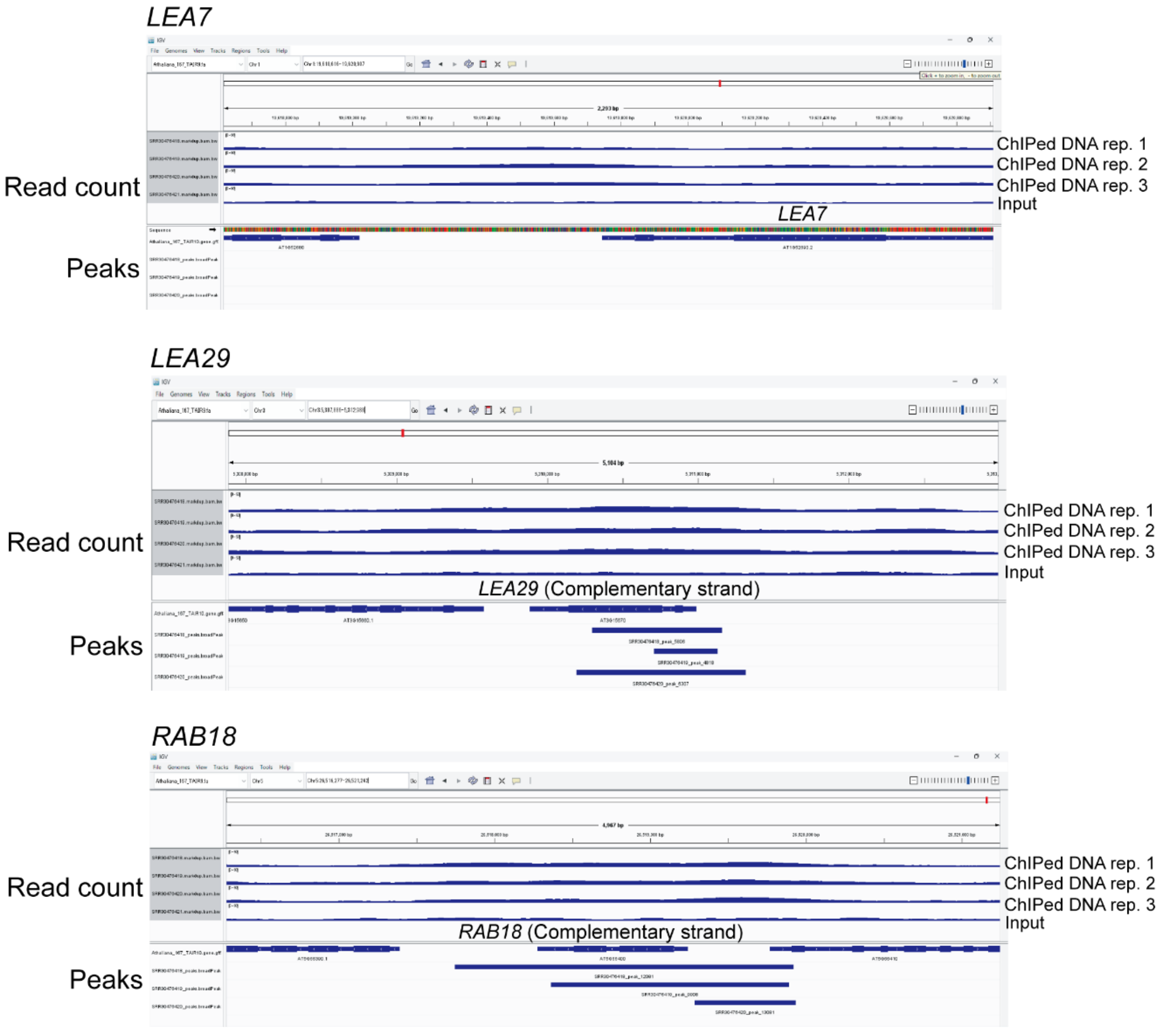


**Fig. S9.** Visualization of ChIP-seq data of the HDA19-FLAG deposited in the Gene Expression Omnibus (GEO) database with accession code GSE275989 (11). was used to visualize the processed ChIP-Seq data at *LEA7*, *LEA29*, and *RAB18* loci. ChIP-seq reads of HDA19 were aligned to the reference *Arabidopsis thaliana* genome (version TAIR9) using bowtie2 2.4.4, and the peak calling using MACS2 2.2.7.1 on alignment results was performed as described by Mehraj et al. (30)

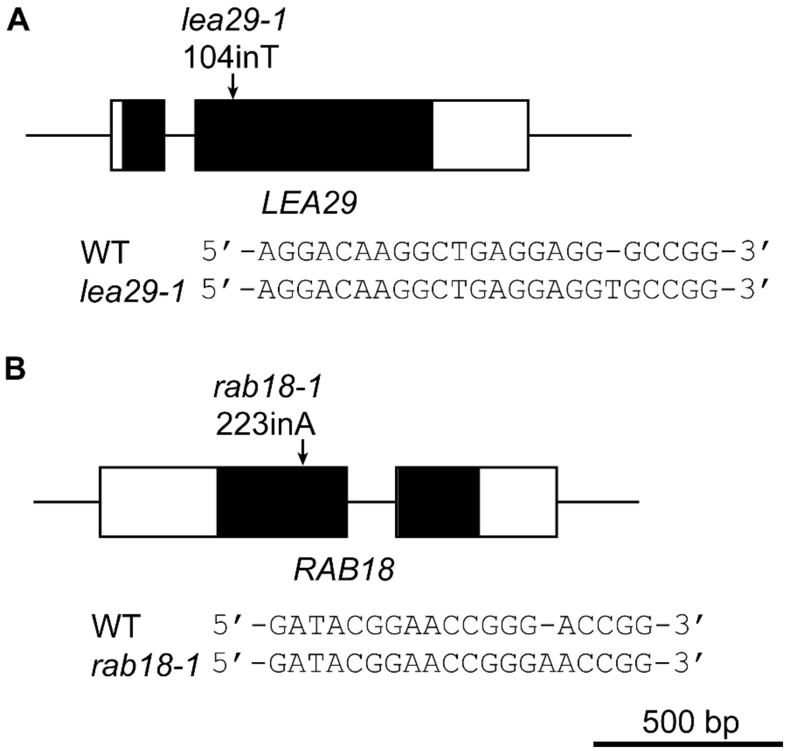


**Fig. S10.** Scheme of the *lea29-1* and *rab18-1* mutations. The two mutants *lea29-1* and *rab18-1* are single nucleotide insertion mutants generated by using clustered regularly interspaced short palindromic repeats (CRISPR)/Cas9 system. (*A*) A thymine insertion occurred 104 nt downstream from the LEA29 translational initiation codon. (*B*) An adenine insertion occurred 223 nt downstream from the RAB18 translational initiation codon.

Fig. 3*C*

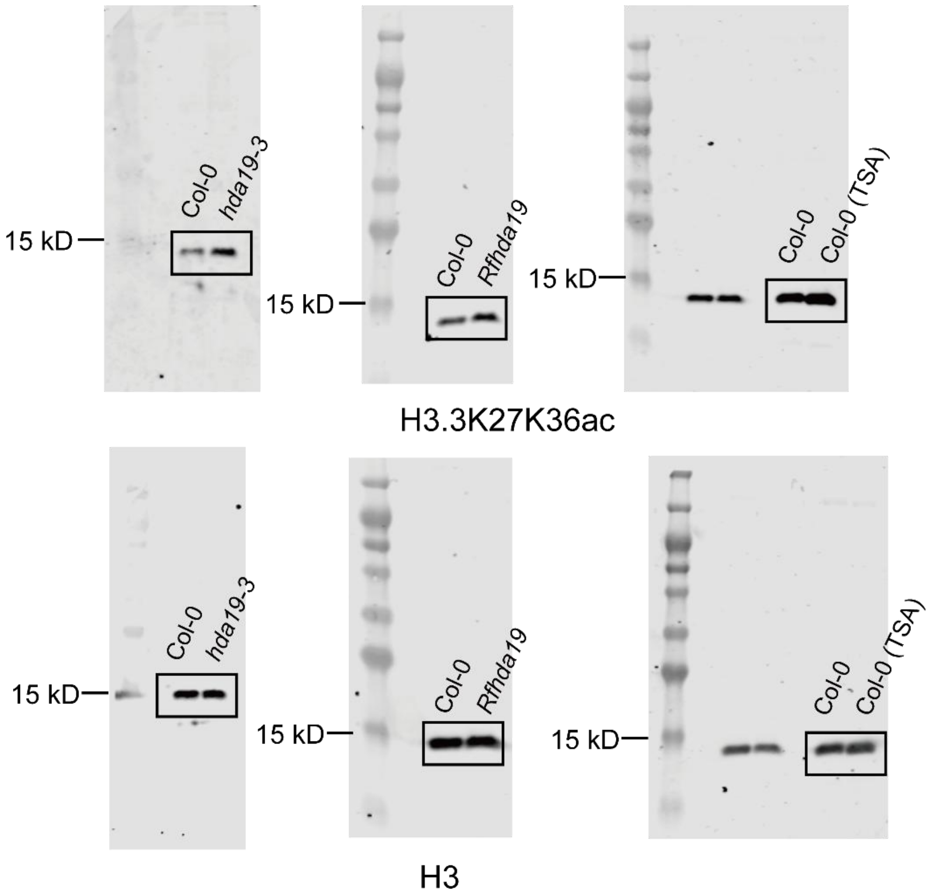


Fig. 3*E*

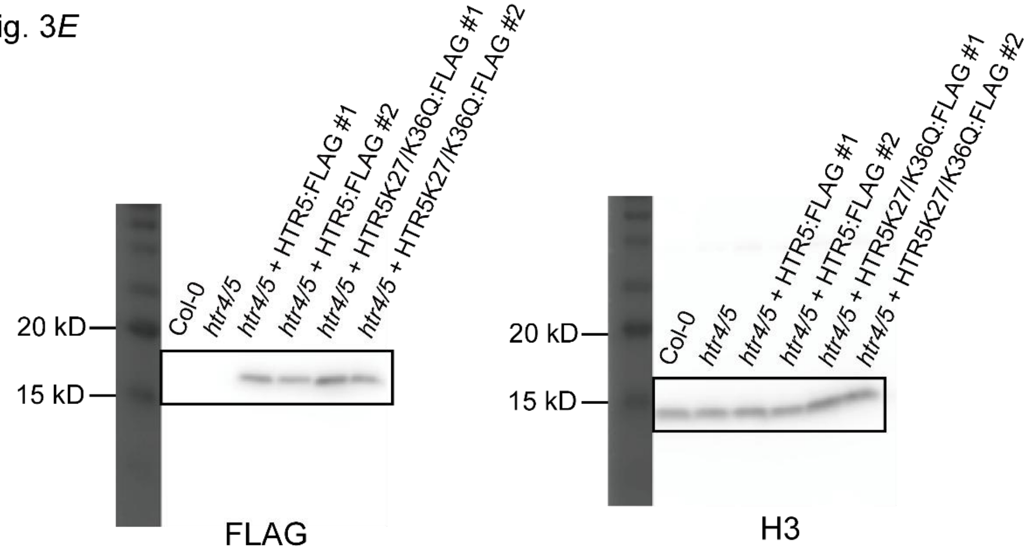


**Fig. S11.** Full-size images of western blots presented in Fig. 3*C* and 3*E*.

**Tables**

Table S1. Identifiers of proteins numbered 1-16 in Fig. 1B and 1C.

| number | symbol/identifier | locus |
|---|---|---|
| **1** | AT2G05580 | AT2G05580 |
| **2** | CRU3 | AT4G28520 |
| **3** | LEA29 | AT3G15670 |
| **4** | PAP85 | AT3G22640 |
| **5** | CRU1 | AT5G44120 |
| **6** | AT4G36700 | AT4G36700 |
| **7** | LEA7 | AT1G52690 |
| **8** | OLEO2 | AT5G40420 |
| **9** | OLEO4 | AT3G27660 |
| **10** | LEA30 | AT3G17520 |
| **11** | CRU4 | AT1G03890 |
| **12** | LEA25 | AT2G42560 |
| **13** | RAB18 | AT5G66400 |
| **14** | OLEO1 | AT4G25140 |
| **15** | LEA48 | AT5G44310 |
| **16** | SESA1 | AT4G27140 |

**Table S2.** H3.1 and H3.3 acK peptides identified with a multiplicity of 2 as shown in Fig. 3A.

| Histone | acK position | Sequence | Mass | Retention time | Score | Delta score |
|---|---|---|---|---|---|---|
| **H3.1 and/or H3.3** | K18, K23 | acKQLATacKAAR | 1069.624 | 54.413 | 204.07 | 117.28 |
| | K18, K23 | acKQLATacKAARK | 1197.719 | 51.049 | 88.596 | 44.053 |
| | K9, K14 | acKSTGGacKAPR | 984.535 | 25.74 | 144.65 | 107.63 |
| **H3.1** | K27, K36 | acK-SAPATGGVacK-KPHR | 1516.847 | 40.87 | 74.853 | 48.729 |
| **H3.3** | K27, K36 | acKSAPTT-GGVacKKPHR | 1546.858 | 41.109 | 118.71 | 81.552 |

**Table S3.** Primers for genome editing.

| Target gene | Forward primer | Reverse primer |
|---|---|---|
| ***RAB18*** | 5’-ATTGAAGGATACGGAACCGGGACC-3’ | 5’-AAACGGTCCCGGTTCCGTATCCTT-3’ |
| ***LEA29*** | 5’-ATTGGGACAAGGCTGAGGAGGGCC-3’ | 5’-AAACGGCCCTCCTCAGCCTTGTCC-3’ |

**Table S4.** Primers for site-direct mutagenesis. Underline shows a mutated codon.

| Mutation site | Forward primer | Reverse primer |
|---|---|---|
| HTR5 K27Q | 5'-GGCTGCACGTcAGTCTGCACC-3' | 5'-GGTGCAGACTgACGTGCAGCC-3' |
| HTR5 K36Q | 5'-TGGAGGAGTCcAGAAGCCCCA-3' | 5'-TGGGGCTTCTgGACTCCTCCA-3' |
| HRT5 full length genome | 5'-CCTTGCCAATCCATATCTGA-3' | 5'-GGGGATGATTGAAGTAGTGG-3' |

**Table S5.** Primers for quantitative RT-PCR.

| Gene | Forward primer | Reverse primer |
|---|---|---|
| *LEA7* | 5’-CACAGAGGAAGTGAAGAGGATAAA-3’ | 5’-CAACAACAAGGATCGGCATAAC-3’ |
| *LEA29* | 5’-GACTTATCAGAGGAAGTGATGAGAA-3’ | 5’-CAGCTGAACCACAACCATAAAG-3’ |
| *RAB18* | 5’-AGAAGAACATGGCGTCTTACC-3’ | 5’-CGGATTTCCGTACTCGTCATAC-3’ |
| *MON1* | 5’-CCGATGACTTGCTTCTACTCTC-3’ | 5’-CTGAGCGTTGTATCTTGGTAGG-3’ |

**Table S6.** Primers for quantitative ChIP-PCR.

| Gene | Forward primer | Reverse primer |
|---|---|---|
| *LEA7* | 5’-TAACACTCGCACATAACTCCAA-3’ | 5’-GTCGGCGAGAGAACAAGATAG-3’ |
| *LEA29* | 5’-AACACACCACGCAGCTATAC-3’ | 5’-CCCAAACCATCGGGACATT-3’ |
| *RAB18* | 5’-CGTCTTACCAGAACCGTCCA-3’ | 5’-CCGTATCCTCCTCCTCCCAT-3’ |

**Table S7. LC-MS/MS data acquisition settings used in this study.**

| Parameter | Proteome *Rfhda19* vs. Wt<br>*hda19-3* vs Wt<br>Acetylome<br>*Rfhda19* vs. Wt | Acetylome *Rfhda19* vs. Wt (unfractionated)<br>*hda19-3* vs. Wt | HDAC pull-down | Myc pull-down | Input Myc pull-down |
|---|---|---|---|---|---|
| **Orbitrap resolution for $MS^1$ scans** | 120,000 | 120,000 | 120,000 | 120,000 | 60,000 |
| **Scan range (m/z)** | 380-1500 | 380-1500 | 280-1500 | 300-1759 | 300-1759 |
| **max. injection time** | 100 ms | 100 ms | 100 ms | 55 ms | 55 ms |
| **automatic gain control target $MS^1$** | 300% | 300% | 300% | 3e6 | 3e6 |
| **Allowed precursor charge state for fragmentation** | 2-6 | 2-6 | 2-6 | 2-5 | 2-5 |
| **Number of dependent scans** | 20 | 20 | 20 | 12 | 15 |
| **Exclusion duration for dynamic exclusion** | 40 sec | 40 sec | 40 sec | 30 sec | 30 sec |
| **Mass tolerance for dynamic exclusion** | ±10 ppm | ±10 ppm | ±10 ppm | ±10 ppm | ±10 ppm |
| **Isolation window (m/z) (no offset)** | 0.7 | 0.7 | 1.6 | 1.2 | 1.3 |
| **Normalized collision energy** | 36% | 36% | 30% | 25% | 25% |
| **Orbitrap resolution for $MS^2$ scans** | 30,000 | 30,000 | 15,000 | 15,000 | 15,000 |
| **Fixed first mass (m/z)** | 100 | 100 | 100 | 100 | 100 |
| **Max injection time** | 86 ms | 150 ms | 150 ms | 150 ms | 55 ms |
| **Automatic gain control target $MS^2$** | 50% | 5% | 5% | 5e4 | 1e5 |

**Dataset S1 (separate file).** Quantitative LC-MS/MS analysis of pull-downs using mini-AsuHd HDAC traps and mini-Lys control probes.

**Dataset S2 (separate file).** MS2-based quantitative analysis of the *Rfhda19* vs WT proteome (*A*) and lysine acetylome (*B*).

**Dataset S3 (separate file).** MS2-based quantitative analysis of the *hda19-3* vs WT proteome (*A*) and lysine acetylome (*B*).

**Dataset S4 (separate file).** Combined analysis of the *hda19* vs WT (*A*) proteome and (*B*) lysine acetylome.

**Dataset S5 (separate file).** Quantitative LC-MS/MS analysis of (*A*) MycHDA19 pull-down and (*B*) input proteome.
